# Catalytic and non-catalytic functions of DNA polymerase kappa in translesion DNA synthesis

**DOI:** 10.1101/2023.11.06.565773

**Authors:** Selene Sellés-Baiget, Sara M. Ambjørn, Alberto Carli, Ivo A. Hendriks, Irene Gallina, Norman E. Davey, Bente Benedict, Bob Meeusen, Emil P.T. Hertz, Laura Slappendel, Daniel Semlow, Sampath A. Gadi, Shana Sturla, Michael L. Nielsen, Jakob Nilsson, Thomas C. R. Miller, Julien P. Duxin

## Abstract

Translesion DNA synthesis (TLS) is an essential process that allows cells to bypass lesions encountered during DNA replication and is emerging as a primary target of chemotherapy. Among vertebrate DNA polymerases, polymerase kappa (Pol() has the unique ability to bypass minor groove DNA adducts in vitro. However, Pol(is also required for cells to overcome major groove DNA adducts but the basis of this requirement is unclear. Here, we combine CRISPR base editor screening technology in human cells with TLS analysis of defined DNA lesions in Xenopus egg extracts to unravel the functions and regulations of Pol(during lesion bypass. Strikingly, we show that Pol(has two main functions during TLS, which are differentially regulated via Rev1 binding. On the one hand, Pol(is essential to replicate across minor groove DNA lesions in a process that depends on PCNA ubiquitylation but is independent of Rev1. On the other hand, via its cooperative interaction with Rev1 and ubiquitylated PCNA, Pol(stabilizes the Rev1-Pol(extension complex on DNA to allow extension past major groove DNA lesions and abasic sites, in a process that is independent of Pol(catalytic activity. Together, our work identifies catalytic and non-catalytic functions of Pol(in TLS and reveals important regulatory mechanisms underlying the unique domain architecture present at the C-terminal end of Y-family TLS polymerases.

## Introduction

Our genome is continuously exposed to endogenous and exogenous DNA damaging agents, which generate lesions that impede DNA synthesis. High fidelity replicative polymerases are unable to accommodate DNA lesions in their catalytic site due to the resultant distortion in DNA geometry (Goodman & Woodgate, 2013; Ling et al., 2001). This often leads to replication fork uncoupling, activation of the S phase replication checkpoint, and replication stress (Cortez, 2019).

DNA lesions encountered during DNA replication can be bypassed by DNA damage tolerance (DDT) mechanisms. To date, two distinct DDT mechanisms have been described: template switching (TS), which relies on a recombination-based process to copy genetic information from the undamaged sister chromatid, and translesion DNA synthesis (TLS), which involves specialized DNA polymerases to synthesize DNA across damaged bases (Friedberg et al., 2002). Unlike replicative polymerases, TLS polymerases exhibit poor processivity and low fidelity. This low fidelity is attributed to the lack of proofreading activity and the presence of a more flexible catalytic site that can accommodate damaged or distorted DNA bases (Goodman & Woodgate, 2013; Ling et al., 2001). Ubiquitylation of the polymerase processivity factor (PCNA) on lysine 164 (K164) plays a crucial role in stimulating both TLS and TS processes (Hoege et al., 2002; Stelter & Ulrich, 2003). This is because most enzymes participating in DDT contain PCNA and ubiquitin binding motifs that bind to ubiquitylated PCNA and are thereby targeted to the lesion site (Sale et al., 2012).

In vertebrates, the primary TLS polymerases operating during DNA replication are Polη, Polι, Polκ, and Rev1, which are classified under the Y-family, and Polζ, which is a member of the B-family. Although a single polymerase may catalyze both steps of DNA lesion bypass – incorporation of a nucleotide opposite the DNA lesion and the subsequent extension past the lesion –TLS often involves the coordinated actions of an “inserter” and an “extender” (Sale et al., 2012). A classic example of an insertion polymerase is Polη, which is specialized in bypassing UV-induced thymine-thymine cyclobutane pyrimidine dimers (CPDs) in an error-free manner (Johnson et al., 1999; Masutani et al., 2000; McCulloch et al., 2004). In contrast, Polζ is a multi-subunit TLS polymerase known for its ability to extend mismatched DNA termini such as the ones originating from nucleotides inserted by Y-family polymerases (Johnson et al., 2000; Lee et al., 2014). This process is observed during the bypass of UV-induced 6-4 photoproducts, cisplatin-induced intrastrand crosslinks, and M.HpaII-based DNA-protein crosslinks (DPCs), where Polζ extends the mispaired nucleotide inserted by Polη (Gallina et al., 2021; Gibbs et al., 2005; Hicks et al., 2010; Yoon et al., 2010).

Despite its deoxycytidyl transferase activity, Rev1 is best known for its scaffolding role for Polζ and other Y-family polymerases (Guo et al., 2003; Martin & Wood, 2019). In fact, Rev1 is an integral component of Polζ both in yeast and in *Xenopus* egg extracts (Acharya et al., 2009; Budzowska et al., 2015). Rev1 also stimulates the recruitment of other Y-family polymerases to lesion sites. This process can occur independently of PCNA ubiquitylation and offers TLS polymerases an alternative recruitment route (Edmunds et al., 2008; Guo et al., 2006). Consistently, Y-family TLS polymerases contain C-terminal regions composed of multiple protein-protein interaction domains that mediate binding to PCNA, ubiquitin, and Rev1. However, the specific interplay between Rev1 and ubiquitylated PCNA in targeting TLS polymerases to lesion sites remains unknown.

In contrast to Polη, Rev1, and Polζ, the role of Polκ in TLS remains poorly understood. Polκ shares homology with bacterial DinB (Gerlach et al., 2001; Ogi et al., 1999). *In vitro,* both DinB and Polκ can bypass adducts located in the minor groove of DNA, which do not fit in the active site of other TLS polymerases (Choi et al., 2006; Jarosz et al., 2006). This is because Polκ contains a unique N-clasp and polymerase associated domain (PAD) that allow its catalytic core to open towards the minor groove of DNA (Jha et al., 2016; Jha & Ling, 2018). Minor groove adducts consist of N^2^-adducted guanosines generated by alkylating agents like benzo[a]pyrene (BP) and acylfuvenes (AF) (Choi et al., 2006; Ogi et al., 2002). Similarly, Illudin S and Mitomycin C (MMC) also generate minor groove DNA lesions, albeit with different chemical properties (Takeiri et al., 2014; Tanasova & Sturla, 2012). Consistently, Polκ deficient cells are exquisitely sensitive to both Illudin S and MMC (Casimir et al., 2023; Olivieri et al., 2020; Williams et al., 2012). However, formal proof that Polκ enables the bypass of minor groove DNA lesions during DNA replication is still lacking.

Intriguingly, Polκ deficient cells are also sensitive to major groove DNA damaging agents, such as UV radiation and cisplatin (Ogi & Lehmann, 2006; Olivieri et al., 2020; Williams et al., 2012), despite Polκ being unable to synthesize across UV lesions *in vitro* (Johnson, et al., 2000). This suggests that Polκ may also have functions in TLS that are independent of its catalytic activity. This was proposed based on the sensitivity of cells expressing a catalytically deficient version of Polκ to different DNA damaging agents (Kanemaru et al., 2015), but the mechanism underlying this non-catalytic function is completely unknown. Moreover, how Polκ is recruited to DNA lesions remains unclear. Like all Y-family polymerases, Polκ contains a long and flexible C-terminal end that mediates interactions with ubiquitylated PCNA. In fact, Polκ contains two PCNA interacting motifs (PIP) and two ubiquitin-binding zinc finger (UBZ) domains that may differently contribute to Polκ targeting to lesion sites. Moreover, if Polκ can be targeted via PCNA-ubiquitylation, why does it also contain a conserved Rev1 interacting region (RIR)? In this regard, Polκ-Rev1 interaction was shown to mediate the formation of a stable Polκ-Rev1-Polζ complex *in vitro* (Xie et al., 2012), but the relevance of this complex during DNA replication is also unknown. In summary, while Polκ catalytic function has been studied *in vitro*, the roles and regulations of Polκ during DNA replication remain elusive, particularly with regards to its function in counteracting major groove DNA lesions.

Here, we elucidate the roles and regulations of Polκ during DNA replication across defined DNA lesions. By combining CRISPR base editor screening technology in human cells with TLS analysis of defined DNA lesions in *Xenopus* egg extracts, we demonstrate that TLS across a minor groove DNA lesion is exclusively dependent on Polκ’s catalytic activity. While this function is dependent on Polκ interaction with ubiquitin and PCNA, it is fully independent of Rev1. In contrast, we find that a Polκ-Rev1 interaction is essential to stimulate the bypass of major groove DNA lesions and abasic sites that require Polζ-mediated extension. Using its flexible C-terminal end which cooperatively binds to Rev1 and ubiquitylated PCNA, Polκ stabilizes the Rev1-Polζ complex on damaged chromatin, allowing extension past DNA lesions in a process that is independent of Polκ catalytic activity. Collectively, our results unravel the multifaceted functions and regulations of Polκ during TLS.

## Results

### Polκ is required to bypass minor groove adducts during DNA replication

Polκ deficient cells are exquisitely sensitive to minor groove inducing agents such as Illudin S (Olivieri et al., 2020). To understand the role of Polκ functional domains in counteracting Illudin S in human cells, we first performed a CRISPR-Cas9 base editor tiling screen in the presence and absence of Illudin S. We designed a single guide RNA (sgRNA) lentiviral library aimed at introducing missense mutations along the Polκ coding sequence. The sgRNA library was cloned into ABE8e-SpG (Sangree et al., 2022), which is a CRISPR base editor system that introduces A◊G conversions in defined proximity to the sgRNA binding site. Using this library, we can target >50% of the amino acids in Polκ with point mutations (**Table S1**). Briefly, RPE1-hTERT p53-/- cells were transduced with the Polκ lentiviral sgRNA library. After selection and expansion, cells were cultured in the presence or absence of a low dose of Illudin S for 12 additional days. Subsequently, sgRNAs were quantified by next generation sequencing to identify editing sites that conferred sensitivity to Illudin S (**Figure 1A** and **Table S1**). Consistent with Polκ unique ability to bypass minor groove DNA adducts *in vitro* (Choi et al., 2006; Jarosz et al., 2006), numerous sgRNAs predicted to cause point mutations in the catalytic domain of Polκ severely sensitized cells to Illudin S without affecting the untreated condition (**Figure 1B**, **S1A**, and **Table S1**). These included several sgRNAs targeting the N-clasp, PAD, palm, and finger domains, which are essential for Polκ TLS properties (**Figure 1B-C** and **S1B-C**) (Boudsocq et al., 2004; Jha et al., 2016; Jha & Ling, 2018). In contrast, no sgRNAs targeting a region of about 80 amino acids within the palm domain sensitized cells to Illudin S (amino acid 210-290; **Figure 1B-C**). This is consistent with the disordered nature of this region where point mutations are unlikely to affect Polκ catalytic function (**Figure S1B-C**). In addition to the catalytic domain, sgRNAs targeting Polκ’s PIP1 (known to bind PCNA) and UBZ2 (known to bind ubiquitin) domains were the only sgRNAs that also significantly sensitized cells to Illudin S (**Figure 1B-C** and **S1B-C**). In contrast, Polκ’s Rev1 interaction region (RIR), UBZ1, or PIP2 did not score, despite the screen containing several sgRNAs that were predicted to introduce missense mutations in each of these domains. These results are consistent with Polκ being targeted to minor groove DNA lesions by binding to ubiquitylated PCNA (via its PIP1 and UBZ2) in a process that is independent of Rev1.

**Figure 1.**
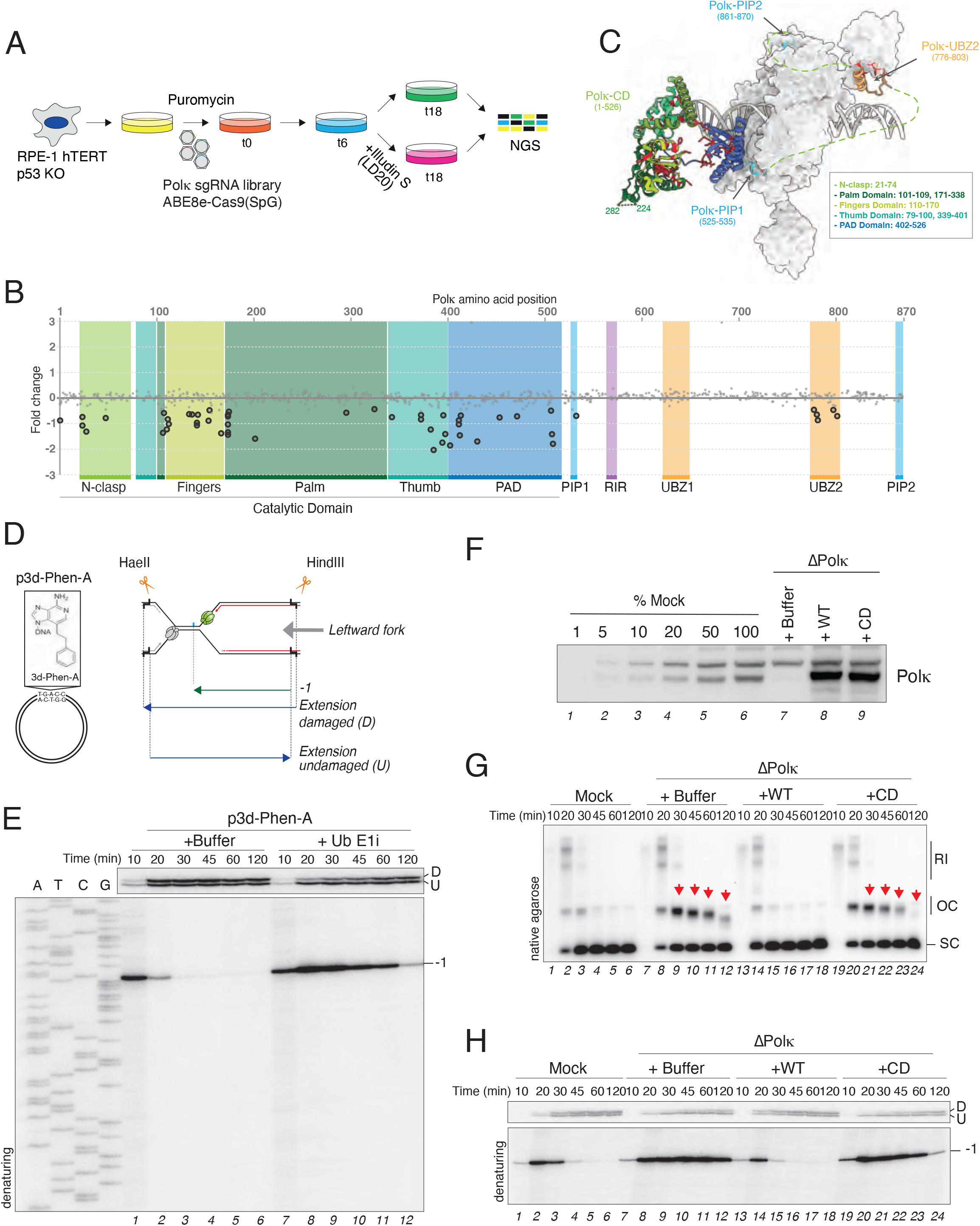
Polκ is essential to bypass minor groove DNA lesions. A) Schematic outline of a Polκ CRISPR base editor tiling screen aimed at identifying missense mutations that sensitizes RPE1-hTERT p53-/- cells to Illudin S. NGS, next generation sequencing. B) Dot plot showing the results of the Polκ CRISPR base editor tiling screen. Each guide is shown as a dot. X-axis represents amino acid position in Polκ, which is targeted for point mutation. Y-axis represents Log2 fold changes between Illudin S and untreated conditions. Larger dots represent guides which are significantly changing in Illudin S (p-value ≤ 0.01). The various domains of Polκ are indicated. C) Composite molecular model of human Polκ with point mutations derived from the base editor tiling screen. Point mutations are highlighted in red. Dashed lines represent disordered regions that are not present in the model. The model was generated by combining structure predictions from AlphaPulldown (Yu et al., 2023) / AlphaFold2 (Jumper et al., 2021), as described in the Methods section, along with deposited structures of human Polκ and PCNA (PDBs: 5W2C (Yockey et al., 2017), 6TNY (Lancey et al., 2020), 3TBL (Zhang et al., 2012)). D) On the left side, schematic of p3d-Phen-A. On the right side, scheme displaying the leftward nascent leading strand and extension products during replication of p3d-Phen-A generated upon HaeII and HindIII digest. Double digestion generates longer damaged and shorter undamaged extension products, which can be resolved on a denaturing polyacrylamide gel. The leftward CMG helicase is depicted in green, while minor groove adduct is depicted in blue. E) p3d-PhenA was replicated in the presence or absence of ubiquitin E1 inhibitor (MLN-7243). Reaction samples were digested with HaeII and HindIII followed by separation on a denaturing polyacrylamide gel alongside a sequencing ladder. The stalling points position in relationship to the minor groove adduct are shown in the lower radiograph. The upper radiograph shows the extension products. Note that the sequencing ladder alignment is shifted 1 nucleotide compared to the nascent strand intermediates. This is due to the faster migration of the sequencing ladder induced by the terminal dideoxynucleotide, as previously shown in Larsen et al., 2019. F) Polκ- depleted extracts were compared to a mock depletion dilution series. Polκ-depleted extracts were supplemented with either buffer (+Buffer), wild-type (+WT), or catalytically inactive (+CD) Polκ. Samples were blotted with Polκ antibody. G) Samples from (F) were used to replicate p3d-Phen-A in the presence of [α-^32^P]dATP. Reaction samples were analyzed by native agarose gel electrophoresis. RI, replication intermediates; OC, open circular; SC, supercoiled. Red arrowheads indicate the accumulation of OC molecules. Note that accumulated OC molecules undergo 5’ to 3’ resection over time, leading to faster migration on the gel. H) Samples from (G) were digested with HaeII and HindIII analyzed as in (E).

To validate our base editor tiling screen and monitor DNA replication across minor groove DNA lesions, we turned to the *Xenopus* egg extract system, which recapitulates both DNA replication and TLS, and thereby allows us to study the bypass mechanism of a single and defined DNA lesion (Gallina et al., 2021; Walter et al., 1998). To this end, we generated a novel plasmid containing a site specific 3-deaza-3-phenethyl-adenosine acylfulvene minor groove DNA adduct (p3d-Phen-A) (**Figure 1D**, left panel) (Malvezzi et al., 2017), and replicated it in egg extracts in the presence of radiolabeled [α-^32^P]dATP. To explore whether the bypass of this lesion requires TLS, p3d-Phen-A and a control plasmid (pCTRL) were first replicated in the presence or absence of a ubiquitin E1 inhibitor, which impedes *de novo* ubiquitylation and TLS (Gallina et al., 2021). The replication reactions were subsequently analyzed using native agarose gel electrophoresis. While addition of the E1 inhibitor did not impact replication of pCTRL (**Figure S1D**, lanes 1-12), it stabilized open circular molecules (OC) up to 120 minutes during replication of p3d-Phen-A (**Figure S1D**, lanes 13-24). This is consistent with a defect in TLS in the absence of *de novo* ubiquitylation, which leads to the accumulation of OC daughter molecules originating from the replication of the adducted parental strand (**Figure S1E**). To understand how minor groove adducts are bypassed, we next analyzed the replication intermediates on a denaturing polyacrylamide gel following the digest with HaeII and HindII (**Figure 1D**, right panel). This allowed us to monitor, at the nucleotide resolution, the leftward nascent leading strand encountering the adduct. During replication of p3d-Phen-A, we observed a specific −1 product (**Figure 1E**, lanes 1-2, bottom radiograph; the 0 position corresponds to the location of the adduct), which was quickly resolved after 20-30 minutes, and correlated with the full extension of the damaged template strand (**Figure 1E**, lanes 2-6; note that ∼70% of the plasmid preparation contains the adduct). This is consistent with nascent strand synthesis first stalling at the adduct and then bypassing the lesion via TLS. In contrast, in the presence of ubiquitin E1 inhibitor, nascent strand synthesis also stalled at −1 but now persisted up to two hours indicating severe TLS inhibition (**Figure 1E**, lanes 7-12).

Next, we investigated whether Polκ is required for bypass of the minor groove adduct. To this end, we depleted Polκ from extracts and replicated p3d-Phen-A (**Figure 1F**). In the absence of Polκ, conversion of OC to supercoiled (SC) molecules was impaired (**Figure 1G**, lanes 7-12, red arrows) and nascent strands persisted at the −1 position (**Figure 1H**, lanes 7-12), indicating the absence of TLS. Polκ depleted extracts were subsequently supplemented with recombinantly purified Polκ wildtype (WT) or a catalytically inactive mutant harboring D199A and E200A substitutions (CD) (**Figure 1F**) (Gerlach et al., 2001), which we confirmed is deficient in DNA synthesis (**Figure S1F**). Importantly, the TLS defect observed upon Polκ depletion was rescued by addition of Polκ WT but not Polκ CD (**Figure 1G-H**, lanes 13-24), confirming that the catalytic activity of Polκ is required for the bypass of minor groove DNA lesions. Thus, *Xenopus egg* extracts are an ideal system to study the functions and regulations of Polκ during TLS.

### Polκ-mediated bypass of minor groove lesions depends on PCNA-ubiquitylation but is independent of Rev1

We next examined the functional domains of Polκ that are required for bypassing minor groove adducts. To this end, we generated different point mutants of Polκ deficient in either PCNA, Rev1, or ubiquitin binding (**Figure 2A**) (Guo et al., 2008; Hishiki et al., 2009; Ogi et al., 2005; Ohashi et al., 2009), which we added back to Polκ depleted extracts. Consistent with our Polκ base editor tiling screen, Polκ depletion was fully rescued by the addition of Polκ RIR* (**Figure 2B**, lanes 13-16, and **S2A-B)**. Moreover, depletion of Rev1 had no impact on the replication of p3d-Phen-A, which was exclusively dependent on Polκ (**Figure 2C**). Thus, Polκ catalytic activity across minor groove DNA lesions is independent of Rev1.

**Figure 2.**
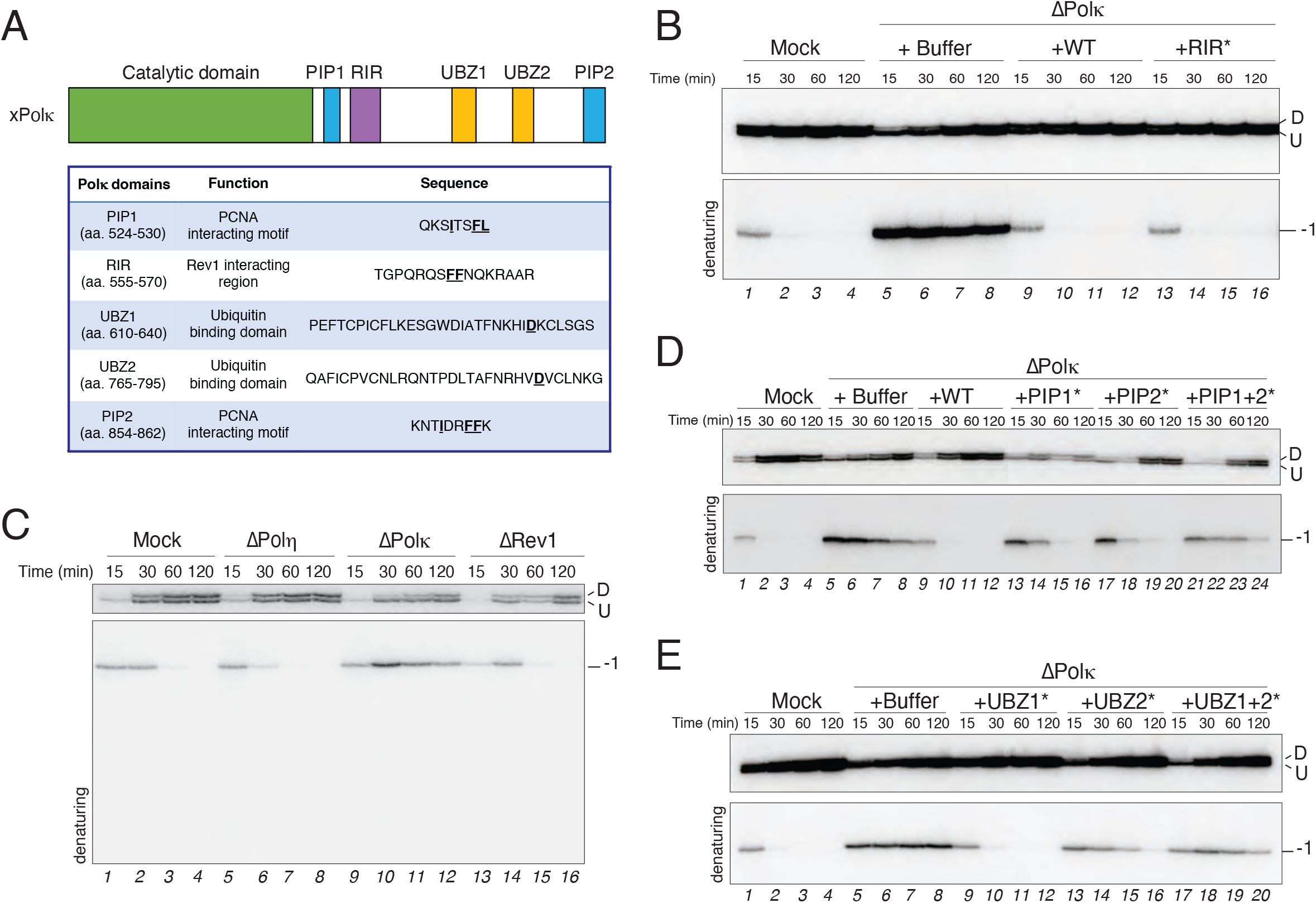
Polκ-mediated bypass of minor groove lesions is independent of Rev1. A) Overview of *Xenopus* Polκ and the specific point mutants used in this study. The underlined amino acids within each domain sequence were mutated to alanine. B) Mock- and Polκ-depleted egg extracts were used to replicate p3d-Phen-A. Polκ-depleted extracts were supplemented with either buffer (+Buffer), recombinant wild-type Polκ (+WT), or Polκ RIR* mutant (+RIR*). Samples were digested and analyzed as in Figure 1E. C) p3d-Phen-A was replicated in mock-, Polη-, Polκ-, or Rev1- depleted extracts. Samples were digested and analyzed as in Figure 1E. D) Mock- and Polκ-depleted egg extracts were supplemented with either buffer (+Buffer), recombinant Polκ wild-type (+WT), PIP1 (+PIP1*), PIP2 (+PIP2*), or PIP1 and PIP2 (+PIP1+2*) Polκ mutants . Extracts were then used to replicate p3d-Phen-A. Samples were digested and analyzed as in Figure 1E. E) Mock- and Polκ-depleted egg extracts were supplemented with either buffer (+Buffer), UBZ1 (+UBZ1*), UBZ2 (+UBZ2*), or UBZ1 and UBZ2 (+UBZ1+2*) Polκ mutants. Extracts were then used to replicate p3d- Phen-A. Samples were digested and analyzed as in Figure 1E.

Analysis of Polκ PCNA interacting motifs revealed that both PIP1 and PIP2 contribute to Polκ-mediated bypass. As seen in **Figure 2D**, addition of either Polκ PIP1* or PIP2* partially rescued the absence of Polκ but not as efficiently as the WT protein (compare lanes 13-16 and 17-20 to 9-12, and **Figure S2C-D**). Moreover, addition of a double PIP mutant (PIP1+2*) further impaired Polκ function (relative to the individual domain mutants) leading to the nascent strand persisting at the −1 position for up to 120 minutes (**Figure 2D**, lanes 21-24, and **S2D**). We conclude from these experiments that both PIP domains can contribute to Polκ TLS activity across minor groove adducts. However, in line with previous work which showed that PIP1 but not PIP2 is required for Polκ mediated synthesis (Lancey et al., 2021; Masuda et al., 2015), and our base editor screen where PIP1 but not PIP2 targeting sensitized cells to Illudin S, we envision that Polκ’s PIP1 is the primary PCNA interactor during catalysis. We hypothesize that PIP2 may act as a recruitment or structural stabilizer by binding to a second PCNA molecule (see discussion).

Lastly, analysis of Polκ UBZ domains further agreed with our base editor screen and revealed that UBZ2 but not UBZ1 is critical for Polκ function (**Figure 2E** and **S2E-F**). Based on these observations, we conclude that Polκ’s catalytic activity across minor groove lesions is independent of Rev1 and may solely depend on PCNA ubiquitylation (see model **Figure 7A** and discussion).

### Polκ-Rev1 interaction is required for the bypass of a major grove DPC

Rev1 is a scaffolding protein that facilitates recruitment of Y-family TLS polymerases to damaged DNA through their RIR domain (Sale et al., 2012). Surprisingly, we find that Polκ’s RIR is dispensable for bypassing minor groove adducts, which raised the question of when the Polκ-Rev1 interaction is needed.

We previously showed that both Polκ and Rev1 are synchronously recruited to DNA during replication of a major groove M.HpaII-DPC containing plasmid (Larsen et al., 2019). DPCs are genotoxic lesions that arise from the covalent linking of proteins to DNA. When a methyl transferase type DPC such as M.HpaII is encountered by the replisome, the protein component of the DPC is first degraded by the SPRTN protease and/or the proteasome, leading to the formation of a DNA-peptide adduct that is then bypassed by TLS polymerases (**Figure S3A** and Duxin et al., 2014; Larsen et al., 2019). TLS across a M.HpaII-DPC is a two-step mutagenic process, in which Polη inserts a nucleotide across the damaged base, followed by Polζ-mediated extension past the peptide adduct (Duxin et al., 2014; Gallina et al., 2021). In the absence of Polη, Polζ performs both insertion and extension but with a higher mutagenesis rate (Gallina et al., 2021). The intriguing recruitment of Polκ to the M.HpaII plasmid with similar kinetics to Rev1-Polζ (**Figure 3A** and Figure 2 in Larsen et al., 2019), raised the possibility that Polκ may also play a function in replication of a major groove DPC lesion.

**Figure 3.**
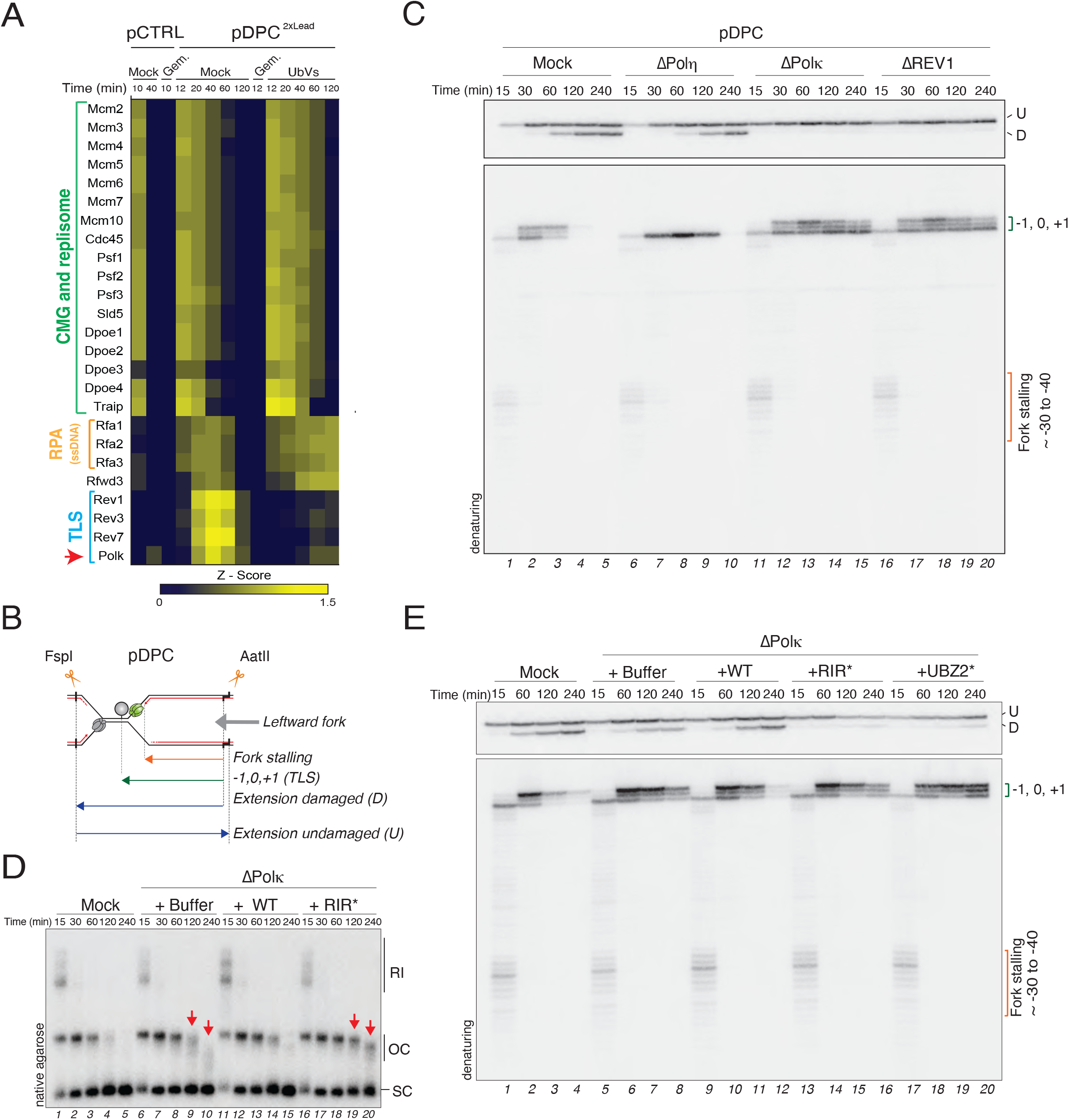
Polκ-Rev1 interaction is required for the bypass of a major groove DPC lesion. A) Heat map displaying the mean of the z-scored Log2 label-free quantification (LFQ) intensity obtained from four biochemical replicates of a control (pCTRL) and a M.HpaII-DPC containing plasmid (pDPC^2xLead^). Geminin was added to block DNA replication where indicated. Ubiquitin-vinyl sulfone (UbVS) was added where indicated to block ubiquitin recycling and deplete the pool of free ubiquitin from the extracts. This data was originally published in Larsen et al., 2019. B) Scheme displaying the nascent leading strand and extension products generated upon FspI and AatII digest of pDPC. Double digestion generates shorter damaged and longer undamaged extension products, which can be resolved on a denaturing polyacrylamide gel. The CMG helicase is depicted in green, while the crosslinked M.HpaII is depicted in grey. C) pDPC was replicated in egg extracts in mock-, Polη-, Polκ- or Rev1-depleted extracts. Reaction samples were digested with FspI and AatII, followed by separation on a denaturing polyacrylamide gel. The stalling points position in relationship to the DPC are shown in the lower radiograph. The upper radiograph shows the extension products. D) Mock- and Polκ-depleted egg extracts were used to replicate pDPC. Polκ-depleted extracts were supplemented with either buffer (+Buffer), wild-type Polκ (+WT), or Polκ RIR mutant (RIR*). Samples were analyzed as in Figure 1G. E) Mock- and Polκ-depleted egg extracts were used to replicate pDPC. Polκ-depleted extracts were supplemented with either buffer (+Buffer), wild-type Polκ (+WT), RIR (+RIR*), or UBZ2 (+UBZ2*) Polκ mutants. Samples were analyzed as in (C).

To investigate whether Polκ is involved in the bypass of DPCs, we first replicated the M.HpaII-DPC plasmid (pDPC) in mock-or Polκ-depleted extracts. As control, pDPC was also replicated in either Polη, or Rev1-depleted extracts (**Figure S3B**; note that Rev1 co-depletes Polζ in egg extracts, thereby enabling only the investigation of the Rev1-Polζ complex) (Budzowska et al., 2015; Gallina et al., 2021). As previously shown, while depletion of Polη had little impact on the conversion of OC to SC molecules (**Figure S3C**, lanes 6-10), depletion of Rev1 stabilized OC molecules for up to 240 minutes, consistent with the absence of bypass across the peptide-DNA adduct (**Figure S3C**, lanes 16-20, red arrows) (Duxin et al., 2014; Gallina et al., 2021). Strikingly, in the absence of Polκ, conversion of OC to SC molecules was also impaired, suggesting a defect in TLS (**Figure S3C**, lanes 11-15, red arrows). To understand how Polκ depletion affects replication across the DPC at the nucleotide resolution, nascent leading strands were analyzed on a denaturing polyacrylamide gel (**Figure 3B**). During replication of pDPC, nascent strand synthesis first stalls ∼30-40 nucleotides upstream of the DPC due to the stalling of the CMG helicase encountering the DPC (**Figure 3C**, lane 1, bottom radiograph). Following CMG bypass of the DPC, the nascent strand advances to the lesion site where it stalls at the −1, 0, and +1 positions (0 refers to the nucleotide where the DPC is crosslinked) (**Figure 3C**, lanes 2-3, bottom radiograph). The nascent strand is then extended past the lesion and fully replicated products appear by 120 and 240 minutes (**Figure 3C**, lanes 3-5, top radiograph). While insertion across the lesion was dependent on Polη (**Figure 3C**, lanes 6-10), extension past the lesion was dependent on Rev1-Polζ (**Figure 3C**, lanes 16-20), as previously described (Gallina et al., 2021). Surprisingly, in the absence of Polκ, nascent strands reached the crosslink with normal kinetics but also permanently stalled at the −1, 0, and +1 positions, mimicking a Rev1-Polζ depletion (**Figures 3C**, lanes 11-15). Importantly, this was not due to co-depletion of Rev1-Polζ with the Polκ antibody (**S3B**, lane 8). Similarly, Polκ was not co-depleted upon Rev1-Polζ depletion (**Figure S3B**, lane 9). Taken together, these results indicate that Polκ plays an essential role in TLS extension past M.HpaII-DPC lesions.

Next, we investigated whether Polκ-Rev1 interaction is needed for this process. To this end, we performed rescue experiments with Polκ WT and Polκ RIR*. While addition of Polκ WT rescued Polκ depletion, addition of Polκ RIR* failed to do so (**Figure 3D-E)**. Thus, in contrast to Polκ-mediated TLS of minor groove adducts, a Polκ-Rev1 interaction is critical for the bypass of a major groove DPC.

### Polκ has a non-catalytic function during DPC bypass

Given that Polκ depletion abolished extension past DPCs, we next investigated whether Polκ polymerase activity stimulates extension. To this end, we performed rescue experiments with Polκ WT and Polκ CD. Strikingly, addition of either Polκ WT or Polκ CD, fully rescued the extension defect caused by Polκ depletion (**Figure 4A-B**), indicating that Polκ has a non-catalytic function in stimulating TLS across the DPC. Consistently, when Rev3, the catalytic subunit of Polζ, was depleted from extracts (**Figure S3B**), extension past DPCs was completely abolished (**Figure S3D**), confirming that Polζ provides the extension activity across the DPC substrate. Identical results were obtained if pDPC was pretreated with proteinase K to digest the protein adduct to a short peptide (Duxin et al., 2014), indicating that the role of Polκ on pDPC replication is independent of DPC proteolysis (**Figure S4A-C**). Moreover, this non-catalytic function of Polκ was also observed when the DPC was placed on either the leading or lagging strand template (**Figure S4D-E**) (Duxin et al., 2014). Together, these results indicate a previously unknown non-catalytic function of Polκ in stimulating Rev1-Polζ mediated extension across a major groove DPC lesion.

**Figure 4.**
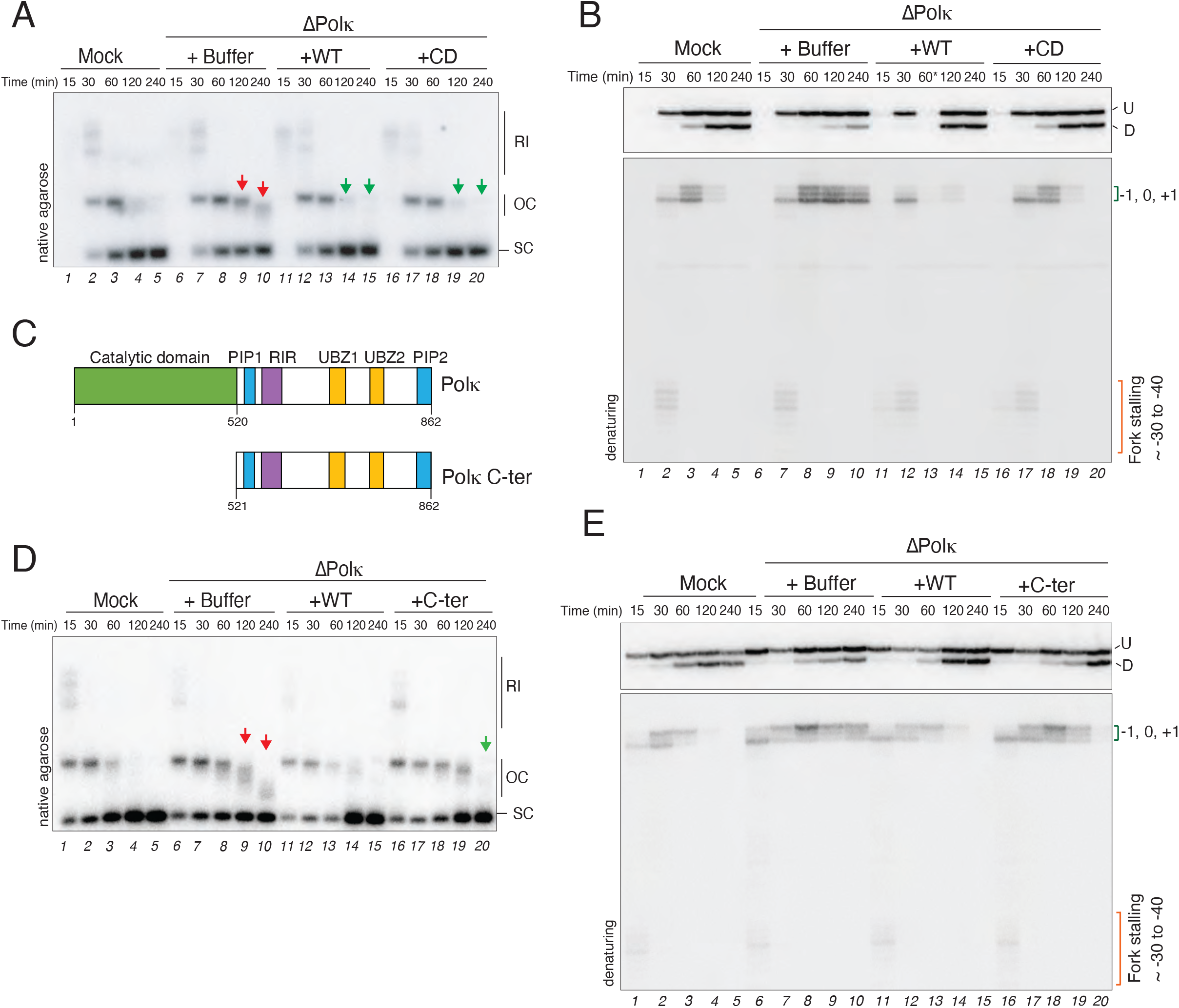
Polκ has a non-catalytic function in stimulating Rev1-Polζ mediated extension across major groove DPC lesions. A) Mock- and Polκ-depleted egg extracts were used to replicate pDPC. Polκ-depleted extracts were supplemented with either buffer (+Buffer), wild-type Polκ (+WT), or catalytically inactive Polκ (+Polκ CD). Samples were analyzed as in Figure 1G. B) Samples from (A) were digested with FspI and AatII and separated on a denaturing polyacrylamide gel. Samples were analyzed as in Figure 3C. * Denotes a sample that was partially lost during DNA extraction (lane 13). C) Schematic of *Xenopus laevis* Polκ showing the domain architecture of full-length (Polκ WT) and C-terminal (C-ter) truncated Polκ proteins. D) Polκ-depleted extracts were supplemented with either buffer (+Buffer), Polκ wild-type (+WT), or Polκ C-terminal fragment (+C-ter). Samples were analyzed as in Figure 1G. E) Samples from (D) were digested with FspI and AatII and separated on a denaturing polyacrylamide gel. Samples were analyzed as in Figure 3C.

To further validate this non-catalytic function, we recombinantly expressed and purified a Polκ C-terminal fragment fully devoid of its N-terminal catalytic domain (Polκ C-ter) (**Figure 4C**). Addition of Polκ C-ter restored extension past the lesion, albeit not as efficiently as addition of the full-length protein (**Figure 4D-E**, compare lanes 11-15 to 16-20). Thus, via its long unstructured C-terminal end, Polκ can stimulate Polζ-mediated extension across a DPC.

Interestingly, addition of Polκ PIP1*, but not PIP2*, restored efficient TLS across DPCs following Polκ depletion (**Figure S4F**). Moreover, addition of Polκ UBZ1*, but not UBZ2*, rescued the TLS defect past the DPC (**Figure S4G** and **3E**). Thus, in contrast to Polκ’s catalytic function across minor groove adducts which is independent of Rev1, Polκ’s non-catalytic function in DPC bypass requires binding to Rev1, but also ubiquitin (via UBZ2), and PCNA (via PIP2) (see model **Figure 7B** and discussion).

### Polκ stimulates Polζ−extension across different DNA lesions

We next investigated whether the non-catalytic function of Polκ is specific to M.HpaII-DPCs or whether it could be extended to other DPCs, such as the ones generated endogenously via HMCES crosslinking (Mohni et al., 2019). It was recently shown that HMCES covalently crosslinks to abasic sites (AP) in ssDNA to protect them from nucleophilic attack and double strand break (DSB) generation (Mohni et al., 2019). Importantly, HMCES-DPCs are endogenous intermediate lesions formed during the repair of abasic site-interstrand crosslinks (AP-ICL) (Semlow et al., 2022). During this process, the NEIL3 glycosylase is recruited to stalled forks and unhooks the AP-ICL by cleaving the N-glycosyl bond of the crosslinked site. This reverses the crosslink and generates an AP site on one of the daughter molecules (**Figure 5A**, i-ii), which is quickly protected by HMCES (Semlow et al., 2016, 2022). HMCES-DPCs are then degraded by the protease SPRTN, and the resulting peptide-DNA adduct is bypassed by Rev1-Polζ (**Figure 5A**, iii) (Semlow et al., 2016, 2022).

**Figure 5.**
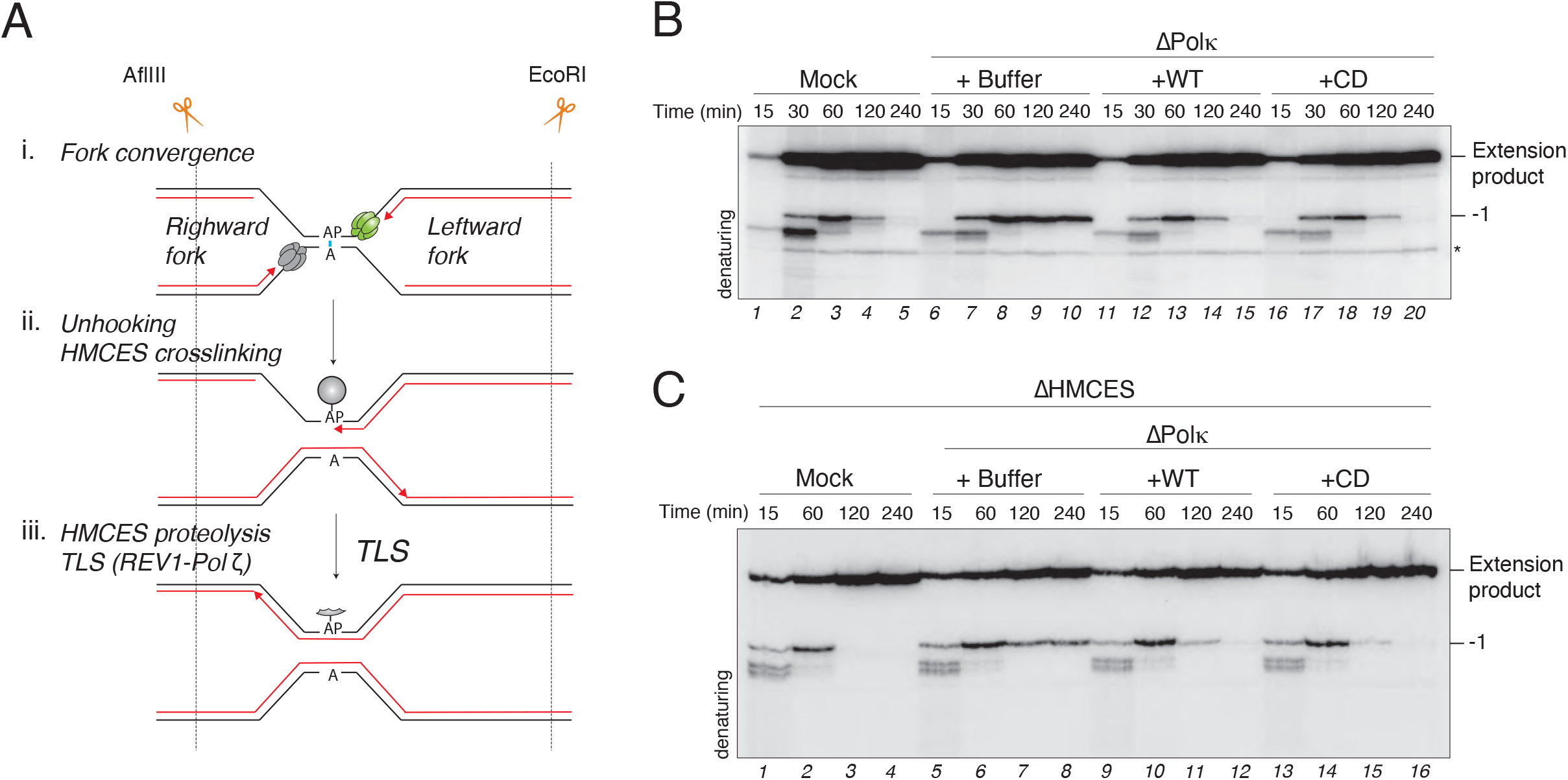
Polκ non-catalytic function stimulates HMCES-DPCs and abasic sites bypass. A) Schematic of HMCES-DPC formation and proteolysis during replication-coupled AP-ICL repair (Semlow et al., 2016, 2022). AflIII and EcoRI allow visualization of stalling points positions relative to the AP-ICL site of the leftward moving fork. B) Mock- and Polκ-depleted egg extracts were used to replicate pAP-ICL. Polκ-depleted extracts were supplemented with either buffer (+Buffer), wild-type (+WT), or catalytically inactive (+CD) Polκ. Samples were subsequently digested with AflIII and EcoRI, followed by separation on a denaturing polyacrylamide gel. Extension products and stalling points position in relationship to the AP-ICL are indicated on the right side of the gel. Asterisk indicates a non-specific digestion product. C) HMCES-depleted extracts were further mock- or Polκ-depleted and used to replicate pAP-ICL. Polκ- depleted extracts were supplemented with either buffer (+Buffer), wild-type (+WT), or catalytically inactive (+CD) Polκ. The samples were analyzed as in (B).

To address whether Polκ also assists Polζ during the bypass of HMCES DPCs, we replicated the AP-ICL containing plasmid in mock- or Polκ-depleted extracts. In the absence of Polκ, permanent stalling at the −1 position was observed for the leftward moving fork, indicating that Polκ is required to bypass HMCES-peptide DNA adducts (**Figure 5B**, compare lanes 1-5 to 6-10, and **S5A**). This effect was rescued by addition of both Polκ WT and Polκ CD (**Figure 5B**, lanes 11-20, **and S5A**). Thus, Polκ non-catalytic function is not restricted to the bypass of M.HpaII-DPCs, but it can be extended to other kinds of DPCs, such as endogenous DPCs generated via HMCES crosslinking to abasic sites.

Next, we addressed whether Polκ is also able to assist Rev1-Polζ across another type of DNA lesion. Abasic sites (AP) are intermediate lesions of AP-ICL repair and dependent on Polζ for their bypass (Haracska et al., 2001). Thus, to stabilize the AP site, we replicated the AP-ICL plasmid in the absence of HMCES (**Figure S5B**) (Semlow et al., 2022). In this setting, Polκ depletion again resulted in the permanent stalling at the −1 position (**Figure 5C**, lanes 5-8, and **S5C**), which was again rescued by the addition of both Polκ WT and Polκ CD (**Figure 5C**, lanes 9-16, and **S5C**). Thus, the non-catalytic function of Polκ extends beyond DPCs to other types of Rev1-Polζ dependent DNA lesions such as AP sites.

### Polκ and Rev1-Polζ form a complex on damaged chromatin

Next, we explored the mechanism by which Polκ promotes Polζ mediated extension. We first hypothesized that Polκ may promote polymerase switching between Polη and Polζ by stimulating the removal of Polη. To test this idea, we monitored Polκ function in TLS in the absence of Polη. Unlike Polη depletion, which is permissive to Polζ mediated bypass (**Figure 6A**, lanes 6-10, and **S6A**) (Gallina et al., 2021), the absence of both Polη and Polκ inhibited TLS across the lesions and nascent strands, now permanently stalled at −1 (**Figure 6A**, lanes 11-15). This was rescued with either Polκ WT or Polκ CD (**Figure 6A**, lanes 16-25, and **S6A**), indicating that Polκ assists Polζ catalysis, even in the absence of Polη. These results suggest that Polκ directly stimulates Polζ recruitment and/or activity during TLS and that this function is independent of the engagement of other TLS polymerases at the lesion site.

**Figure 6.**
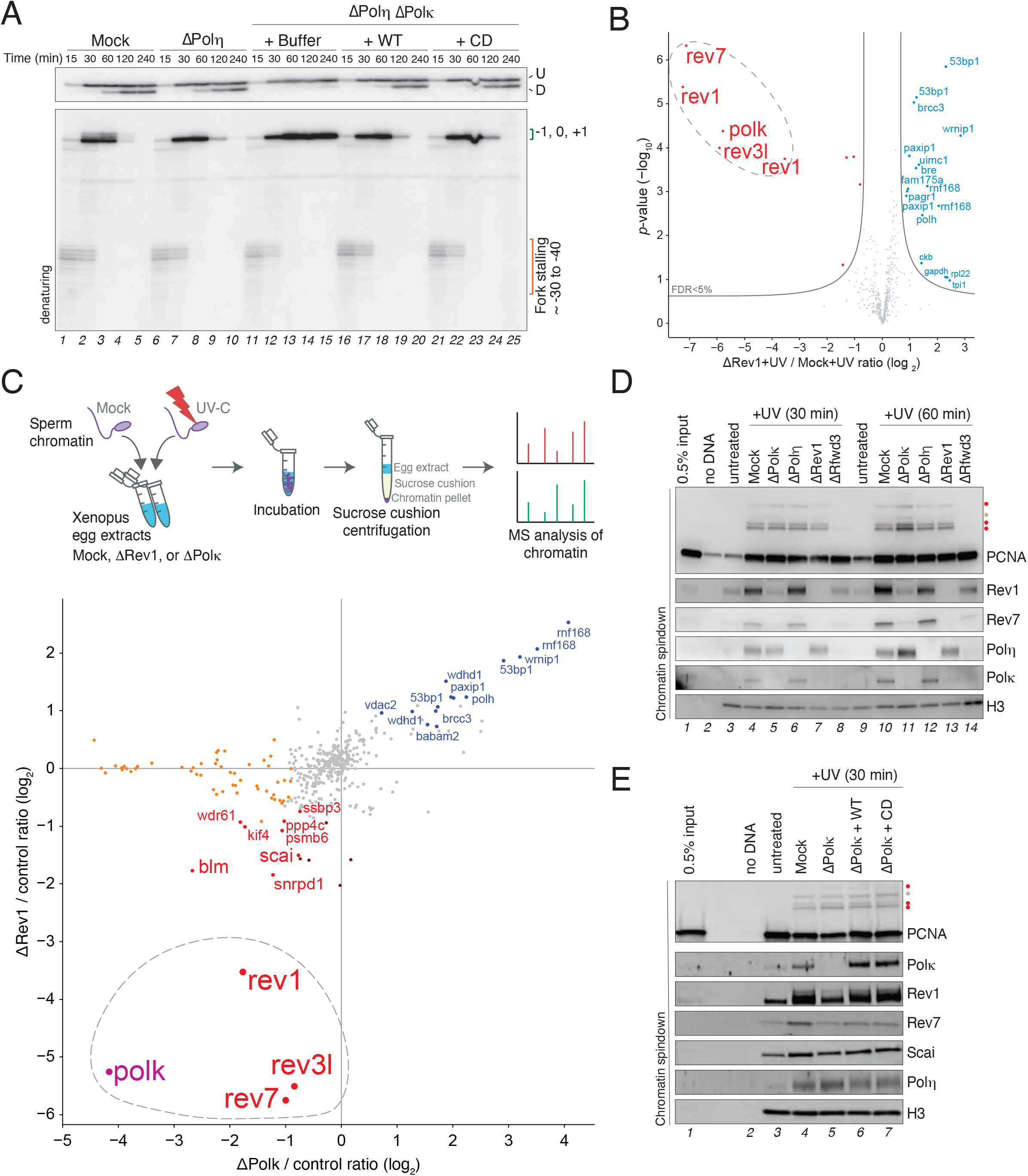
Polκ and Rev1-Polζ form a stable complex on damaged DNA. A) Mock-, Polη-, or Polη- and Polκ-depleted egg extracts were used to replicate pDPC. Polκ- depleted extracts were supplemented with either buffer (+Buffer), wild-type (+ WT), or catalytically inactive (+CD) Polκ. Samples were subsequently digested with FspI and AatII and analyzed as in Figure 3C. B) Chromatin mass spectrometry (CHROMASS) analysis of protein recruitment to UV-treated sperm chromatin in mock- or Rev1- depleted extracts. The volcano plot illustrates the difference in abundance of proteins between the two sample conditions (x axis), plotted against the p-value resulting from two-tailed Student’s two-sample t-testing (y axis). Proteins significantly down- or upregulated (false discovery rate [FDR] < 5%) in Rev1-depleted reactions are represented in red or blue, respectively. n = 4 biochemical replicates. FDR < 5% corresponds to permutation-based, FDR-adjusted p-value (i.e. q-value) < 0.05. Different isoforms of the same protein can be detected (e.g., Rev1) (data originally published in Gallina et al., 2021). C) Schematic of CHROMASS (top panel). Graph illustrating protein recruitment to UV-treated sperm chromatin in the presence or absence of Rev1 or Polκ, as determined by CHROMASS analysis (Table S2). Red dots (bottom-left quadrant) indicate the proteins that were significantly de-enriched on sperm chromatin both in the absence of Rev1 and Polκ. Blue dots (top-right quadrant) indicate the proteins that were significantly enriched on sperm chromatin both in the absence of Rev1 and Polκ. Orange dots indicate the proteins that were significantly de-enriched on sperm chromatin in the absence of Polκ. n=4 biochemical and n=8 technical replicates, significance was determined via two-tailed Student’s two-sample t-testing, with permutation-based FDR control (s0=0.5) to ensure an adjusted p-value (i.e. q-value) of <0.05. Note that different isoforms of the same protein can sometimes be detected. D) Sperm chromatin was either untreated or treated with 2000 J/m2 of UV-C and then added to non-replicating mock-, Polη-, Polκ-, Rev1- or Rfwd3-depleted extracts. Chromatin was isolated, and the proteins associated were blotted with the indicated antibodies. Red dots correspond to PCNA ubiquitylation (mono-, di-, and tri- ubiquitin); the grey dot corresponds to mono-sumoylated PCNA (Gallina et al., 2021). E) Sperm chromatin was either untreated or treated with 2000 J/m2 of UV-C and added to non-replicating Polκ-depleted extracts. Polκ-depleted extracts were supplemented with either buffer (+Buffer), wild-type (+WT), or recombinant catalytically inactive Polκ (+CD). Chromatin was isolated, and the proteins associated were blotted with the indicated antibodies as in (D).

**Figure 7.**
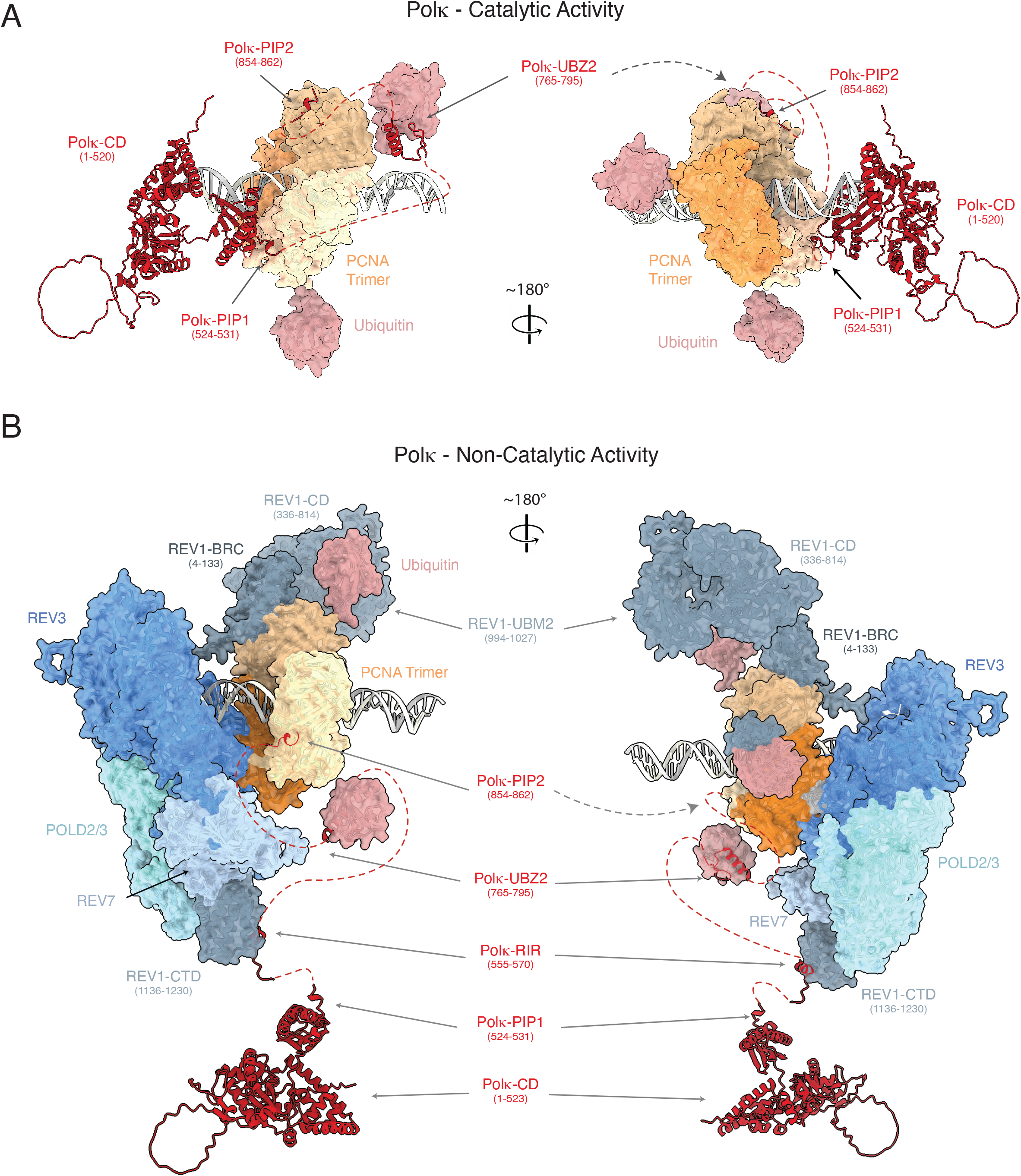
Molecular models of Polκ catalytic and non-catalytic functions during TLS. Composite molecular models of the catalytic (A) and non-catalytic (B) functions of Polκ during TLS. Models were generated by combining structure predictions from AlphaPulldown (Yu et al., 2023) / AlphaFold2 (Jumper et al., 2021) as described in Methods. A) During its catalytic function, Polκ can interact with PCNA monomers via its PIP1 or PIP2 domains bringing its catalytic domain into close proximity with the DNA substrate. The long flexible Polκ C-terminal region additionally allows Polκ to interact with a monoubiquitinated PCNA through its UBZ2 domain. B) During its non- catalytic function, Polκ can bind Rev1 through its RIR domain; the UBZ2 and PIP2 domains are able to bind to monoubiquitinated-PCNA thanks to the long flexible Polκ C-terminal region. Notably, whilst the Polκ RIR domain is Rev1-bound, the PIP1 domain is positioned such that it would be unable to interact with monoubiquitylated PCNA.

We have previously shown that upon UV damage, Polκ is de-enriched on damaged chromatin when Rev1-Polζ is depleted from extracts (**Figure 6B** and Gallina et al., 2021). Since Polκ and Rev1 do not co-deplete each other (**Figure S3B**), this suggests that Polκ may form a complex with Rev1-Polζ on damaged chromatin. To test the potential interdependency between Polκ and Rev1-Polζ localization to damaged chromatin, we performed chromatin mass spectrometry (CHROMASS) to identify proteins whose recruitment depend on either Polκ or Rev1 (Räschle et al., 2015). Briefly, sperm chromatin was treated with a high dose of UV-C, followed by its incubation in either mock-, Polκ-, or Rev1-depleted egg extracts, and analysis by quantitative high-resolution mass spectrometry (**Figure 6C**, top panel). In agreement with the idea that Rev1-Polζ and Polκ form a complex on damaged DNA, Rev1 depletion led to the significant de-enrichment of Polκ from damaged chromatin (**Figure 6C**, Y axis, **S6B-C,** and **Table S2**). Similarly, Polκ depletion also led to the de-enrichment of Rev1 and Polζ (Rev3, and Rev7) (**Figure 6C**, X axis, **S6D**, and **Table S2**). Note that Scai, which associates with Rev1-Polζ (Schubert et al., 2022), was also significantly de-enriched in both conditions (**Figure 6C**). In addition to Polζ, we also noted that many Fanconi Anemia (FA) proteins were significantly de-enriched from Polκ depleted samples (**Figure S6D**, and **Table S2**). This was likely caused by a co-depletion of the FA complex with the Polκ antibody (**Figure S6E**). Conversely, upon depletion of either Rev1 or Polκ, we observed the enrichment of Polη and its interactor Wrnip1 (Yoshimura et al., 2014; Yuasa et al., 2006), as well as proteins involved in double-strand break (DSB) repair, such as Rnf168 and 53bp1 (**Figure 6C**, and **Table S2**). This suggests that in the absence of either Polκ or Rev1-Polζ, gap-filling synthesis across certain UV lesions is disrupted (e.g. UV induced 6-4 photoproducts), which leads to the accumulation of Polη on chromatin and ultimately triggers the formation of DNA DSBs. Alternatively, the accumulation of 53bp1 could be linked to the formation of large ssDNA gaps that form when TLS is inhibited and to the recently described role of 53bp1 in regulating the balance between TLS and TS (Chen et al., 2022).

To validate our CHROMASS results and the formation of a specific Rev1-Polζ-Polκ complex on damaged DNA, we assessed protein recruitment to UV damaged sperm chromatin by immunoblotting upon Rev1 or Polκ depletion. As controls, we also depleted Polη or Rfwd3, which regulates PCNA ubiquitylation and TLS polymerase recruitment to lesion sites (Gallina et al., 2021; Hoege et al., 2002; Stelter & Ulrich, 2003). Consistently, Rfwd3 depletion led to impaired PCNA ubiquitylation and de-enrichment of Polη, Rev1-Polζ, and Polκ to UV treated chromatin compared to the control reaction (**Figure 6D**, compare lanes 4 to 8 and 10 to 14). Confirming the CHROMASS results, the enrichment of Rev1-Polζ was also impaired in the absence of Polκ (**Figure 6D**, compare lanes 4 to 5 and 10 to 11). Similarly, the recruitment of Polκ was abolished in the absence of Rev1 (**Figure 6D**, compare lanes 4 to 7 and 10 to 13). In contrast, Polη enrichment to damaged chromatin was independent of either Rev1 or Polκ and conversely, the absence of Polη did not impact Rev1-Polζ or Polκ recruitment (**Figure 6D**). The loss of Rev1-Polζ recruitment observed upon Polκ depletion could be rescued by addition of either Polκ WT or Polκ CD (**Figure 6E**, compare lanes 5 to 6-7). Together, these results indicate that Polκ and Rev1-Polζ form a stable complex on damaged DNA, which is independent of other Y-family TLS polymerases such as Polη.

## Discussion

Polκ has the unique ability among eukaryotic polymerases to bypass minor groove DNA adducts (Choi et al., 2006; Jarosz et al., 2006). Here, we study for the first time the bypass mechanism of minor groove DNA lesions at nucleotide resolution using a whole extract system that recapitulates DNA replication and TLS. We demonstrate that bypass of a minor groove DNA lesion is exquisitely dependent on Polκ catalytic activity, validating previous observations made with purified proteins. Strikingly, we also uncover a non-catalytic function of Polκ during TLS that is required for Polζ- mediated extension past abasic sites and major groove DNA lesions such as DPCs. Importantly, Polκ’s non-catalytic function in TLS requires an interaction with Rev1, while Polκ catalytic function is Rev1-independent. Thus, Polκ possesses multiple functions during TLS that are differently regulated via its C-terminal unstructured end, implications of which we discuss below.

### Polκ and minor groove DNA lesions

Previous *in vitro* work showed that Polκ is the only vertebrate DNA polymerase that can synthesize across minor groove DNA lesions, but direct evidence of this process in a physiological context was lacking (Choi et al., 2006; Jarosz et al., 2006). Using *Xenopus* egg extracts, we show that Polκ is essential to bypass minor groove DNA adducts (**Figure 1**). In the absence of Polκ, nascent strands remained stalled at the lesion for up to 2 hours (**Figure 1H**), indicating that no other DNA polymerase can compensate for Polκ deficiency. This is consistent with the structural features of Polκ, where the unique conformation of the PAD opens the catalytic core towards the minor groove of DNA (Jha et al., 2016). Moreover, Polκ contains a unique N-terminal region known as the N-clasp, which also contributes to stabilizing the incoming template DNA (Jha et al., 2016). In accordance with this, our base editor screen suggests that point mutations in several amino acids in the PAD and N-clasp domains, increase cellular sensitivity to Illudin S when mutated (**Figure 1B**).

We further provide the regulatory mechanism underlying Polκ’s catalytic function. Previous *in vitro* studies reported that Polκ’s interaction with PCNA through its PIP1 domain is required for Polκ’s catalytic activity, whereas the second PIP2 domain is dispensable (Lancey et al., 2021; Masuda et al., 2015). Our base editor screen showed that targeting PIP1 but not PIP2 increased the sensitivity to Illudin S, further supporting the notion that PIP1 mediates the primary Polκ-PCNA interaction needed for DNA synthesis (**Figure 1B**). However, our work in *Xenopus* egg extracts suggests that PIP2 can also contribute to Polκ function. We show that while either mutation of PIP1 or PIP2 mildly affected Polκ catalytic function, mutation of both PIPs severely impaired lesion bypass (**Figure 2D**). This suggests that while Polκ binding to PCNA is essential, either PIP could partially compensate for each other during catalysis. Consistent with our data, a composite molecular model of the catalytic complex suggests that both PIP domains are capable of binding DNA-loaded PCNA whilst Polκ synthesizes DNA (**Figure 7A**). In fact, each PIP could in principle bind to a different PCNA molecule of the PCNA trimer, thereby further stabilizing Polκ on the damaged template (**Figure 7A**). In this scenario, we envision that each PIP binds successively, where PIP2 first binds a free PCNA molecule of the PCNA trimer before PIP1 binds to a second PCNA molecule and thereby locks the catalytic site on the damaged template.

In addition to PCNA binding domains, Polκ also contains a Rev1 interacting region. Rev1 has been shown to act as a scaffolding protein that recruits Y-family TLS polymerases to damaged sites (Guo et al., 2003). However, we show that Polκ-dependent bypass of minor groove adducts does not require Rev1 nor Rev1 binding (**Figure 2B-C**). Instead, Polκ’s interaction with ubiquitylated PCNA (via PIP1, PIP2, and UBZ2), is sufficient for Polκ-mediated synthesis (**Figure 7A**). Our findings enable reinterpretation of a previous study of Polκ function during replication-independent ICL repair, which was dependent on Polκ catalytic activity but independent of its interaction with Rev1 (Williams et al., 2012). In this study, the authors investigated the repair mechanism of an ICL lesion reminiscent of ICLs induced by MMC, which result in a minor groove DNA adduct following a first round of nucleotide excision by nucleotide excision repair (NER). In light of our findings, these results are likely due to the use of a minor groove DNA lesion, which requires Polκ catalytic function and is therefore independent of Rev1, rather than to a direct regulation of Polκ by NER during replication-independent ICL repair.

### Polκ non-catalytic function in TLS

If Polκ catalytic function is independent of Rev1, when does Polκ-Rev1 interaction become relevant? We show that Polκ, via its interaction with Rev1, stimulates Polζ-mediated extension across different DNA lesions (**Figure 3-5**). Strikingly, this stimulatory effect is fully independent of Polκ’s catalytic activity (**Figure 4**). In fact, a truncated Polκ devoid of its catalytic domain was still able to stimulate Polζ-mediated bypass, albeit not as efficiently as the full-length protein (**Figure 4D-E**). This suggests that most of the function needed to stimulate Polζ-mediated extension is driven by Polκ C-terminal domains, which interact with ubiquitylated PCNA and Rev1 (**Figure 7B**) (Vaisman & Woodgate, 2017). The lower efficiency of rescue by the Polκ C- terminal fragment could be attributed to its highly disordered structure, which may be less stable than the full-length protein in solution.

Through the superimposition of AlphaFold2 generated molecular models of *Xenopus* Rev1-Polζ onto the known Rev1-Polζ structure from yeast (Malik et al., 2020), and modeling of the interactions between Polκ and this complex, we find that the Polκ- Rev1 interaction via the RIR domain is likely to be mutually exclusive with a PIP1 interaction with PCNA (**Figure 7B**). Consistently, Polκ non-catalytic function requires Polκ RIR and PIP2 but is independent of PIP1 (**Figure 3D-E** and **S4F-G**). Based on our findings that: (1) Rev1-Polζ-Polκ forms a stable complex on damaged chromatin, and (2) the formation of this complex is reciprocally-dependent on both Polκ and Rev1- Polζ (but independent of other Y-family polymerases; **Figure 6**), we propose that Polκ utilises its multiple interaction domains to stabilize the Rev1-Polζ complex on DNA, promoting Polζ-mediated extension beyond different DNA lesions (**Figure 7B**). This may enable the Rev1-Polζ-Polκ complex to act as the last resort TLS complex, only functioning following failed attempts by Y-family polymerases to bypass a lesion (i.e. Polη, Polκ, Rev1, or Polι). These would include lesions across which Y-family polymerases can insert but not extend (e.g. 6-4 photoproducts or M.HpaII-DPCs) but also lesions that are unsuited to Y-family polymerase insertion (e.g. M.HpaII-DPCs in the absence of Polη; **Figure 6A**). Notably, our results advance the findings of a previous structural study that showed the formation of a stable Rev1, Rev3, Rev7, and Polκ quaternary complex *in vitro* (Xie et al., 2012) by now demonstrating the functional relevance of this complex.

Interestingly, in addition to its catalytic function across minor groove DNA lesions, Polκ also possesses extension activity past mismatched bases *in vitro* (Haracska et al., 2002; Washington et al., 2002). The formation of this Rev1-Polζ-Polκ complex enables the recruitment of two distinct polymerases with extension activity to the same DNA lesion. Although we only observe the stimulation of Polζ by Polκ, it is tempting to speculate that the opposite might also occur on certain DNA lesions that Polζ may not be able to extend. This arrangement would provide a safeguarding mechanism, ensuring that if one polymerase is unable to extend past the DNA lesion, the other polymerase could quickly replace it and extend beyond the lesion. Together, our data reveal both catalytic and non-catalytic functions of Polκ during TLS and demonstrate the specific roles of its different interaction domains for each function.

### Role of Rev1 in Y-family TLS regulation

Y-family TLS polymerases contain long flexible C-terminal ends composed of multiple interaction domains with PCNA, ubiquitin, and Rev1. The existing models propose that PCNA ubiquitylation facilitates the recruitment of TLS polymerases to sites of DNA lesions (Hoege et al., 2002; Stelter & Ulrich, 2003). Alternatively, Rev1 acts as a scaffold that can recruit TLS polymerases to damaged sites in the presence or absence of PCNA ubiquitylation (Guo et al., 2003, 2006). Thus, two modes of TLS polymerase recruitment co-exist but the interplay and conditions under which one or the other pathway becomes relevant is unknown. In our study, we show that binding of Polκ to ubiquitylated PCNA is essential to target its catalytic function in a process independent of Rev1 (**Figure 2**). In contrast, Rev1 regulates Polκ non-catalytic function in stimulating Polζ-extension (**Figure 3D-E**). Thus, rather than acting as a general recruitment platform for Y-family polymerases, Rev1 appears to regulate the formation of specialized TLS sub-complexes. Like Polκ, Polη and Polι also contain RIR domains of unclear significance and it is tempting to speculate that Rev1 might also regulate unknown functions of these polymerases.

### Polκ function in mammalian cells

How do our findings in *Xenopus* egg extracts clarify observations made in mammalian cells? Consistent with Polκ essential function to replicate across minor groove DNA lesions, Polκ knockout cells exhibit severe sensitivity to minor groove inducing agents such as Illudin S and MMC (Olivieri et al., 2020; Williams et al., 2012), which we confirmed here via our Polκ base editor screen. Intriguingly, Polκ knockout cells are also sensitive to other DNA damaging agents, such as UV and cisplatin, which induce lesions on the major groove of DNA that require Polζ-mediated extension (Budzowska et al., 2015; Martin & Wood, 2019; Ogi & Lehmann, 2006; Olivieri et al., 2020). Hence, in light of our discoveries, we propose that the sensitivity of Polκ knockout cells to major groove inducing agents like UV and cisplatin is attributed to the non-catalytic function of Polκ, facilitating Rev1-Polζ-dependent bypass of these lesions. Accordingly, previous evidence suggested a non-catalytic function of Polκ in replication fork restart or counteracting the sensitivity to different DNA damaging agents (Kanemaru et al., 2015; Tonzi et al., 2018). Moreover, Polκ knockout cells are also sensitive to oxidative agents, such as potassium bromate, which generates abasic sites and HMCES-DPCs (Olivieri et al., 2020). This sensitivity is also likely to be attributed to the non-catalytic role of Polκ across abasic sites or HMCES-DPCs, which we report here (**Figure 5**).

In contrast to Polκ knockouts, Rev3–/– mouse embryonic stem cells are not viable (Lange et al., 2012), which suggests that Polζ possesses Polκ independent functions. Interestingly, Rev3 also functions beyond TLS by facilitating DNA replication through heterochromatic regions (Ben Yamin et al., 2021), which could account for its essential role.

Finally, the development of TLS inhibitors is an emerging strategy to enhance tumor sensitivity to first-line chemotherapeutics (Wojtaszek et al., 2019; Yamanaka et al., 2017). Our results highlight how targeting defined functional domains of Y-family polymerases such as the Polκ-Rev1 interaction might selectively sensitize cancer cells to specific genotoxic agents.

## Materials and methods

### *Xenopus* egg extracts and DNA replication reactions

*Xenopus* egg extracts were prepared as described previously (Lebofsky et al., 2009). All experiments involving animals were approved by the Danish Animal Experiments Inspectorate and conform to relevant regulatory standards and European guidelines.

For plasmid DNA replication, plasmids were licensed in high-speed supernatant (HSS) at a final concentration of 7.5 ng/µL for 30 minutes at room temperature (RT). Replication was initiated by adding two volumes of nucleoplasmic egg extract (NPE). Gap filling reactions were performed in non-licensing extracts, were one volume of HSS was premixed with two volumes of NPE prior to the addition of plasmid DNA (final concentration of 10 ng/µL). For replication in the presence of LacI, plasmid DNA (150 ng/µL) was incubated with an equal volume of 12 µM LacI for 1 hr prior to licensing (Duxin et al., 2014). The Ubiquitin E1 inhibitor MLN-7243 (Active Biochem) was supplemented to NPE at a final concentration of 200 µM, 10 minutes before initiating the reaction. To visualize DNA replication intermediates, replication reactions were supplemented with [α-^32^P]dATP (Perkin Elmer). For each timepoint, 1 µL of the reaction mixture was added to 5 µL of stop buffer (5% SDS, 80 mM Tris pH 8.0, 0.13% phosphoric acid, 10% Ficoll), followed by addition of 1 µL of Proteinase K (20 mg/mL) (Roche). The samples were incubated at 37°C for 1 hour and subsequently separated by using 0.9% native agarose gel electrophoresis, and visualization using a phosphorimager. The radioactive signal was quantified using ImageJ (NIH, USA).

### Preparation of DNA constructs

pDPC and pDPC^2xLead^ were previously described in Larsen et al., 2019. Additionally, pDPC^PK^ and pDPC^ssDNA-PK^ were previously described in Gallina et al., 2021 as pMH^PK^ or pMH^ssDNA-PK^, respectively. Moreover, pDPC^Lead^ and pDPC^Lag^ were previously described in Duxin et al., 2014 as pDPC-L^Top^ ^(Lead)^ and pDPC-L^Bot^ ^(Lag)^, respectively. To generate a plasmid containing a 3-deaza-3-phenethyl-adenosine acylfulvene lesion (p3d-Phen-A), we first removed the LacO array from pJSL3 (Sparks et al., 2019) using the complementary overhangs of the BsrG1 and BsiWI restriction sites. Subsequently, the two Nb.BsrdI nicking sites were removed using mutagenesis. An A located in the position 1557 of the plasmid was mutated into a C to remove the first Nb.BsrDI site using the following primers: 5’- CCACGATGCCTGTAGC**C**ATGGCAACAACGTTGC-3’ and 5’-GCAACGTTGTTGCCATGGCTACAGGCATCGTGG-3’. Second, a T located in the position 1740 of the plasmid was mutated into a C to remove the second Nb.BsrDI site using the following primers: 5’-GGTCTCGCGGTATCAT**C**GCAGCACTGGGGCCAG- 3’ and 5’-CTGGCCCCAGTGCTGCGATGATACCGCGAGACC-3’. Afterwards, we used the PciI and BsaX1 restriction sites to clone in the following oligo 5’- CATGGCTCTTCNACCTCAACTACTTGACCCTCCTCATTCATTGCTTG-3’ to introduce Nt.BspQ1 and Nb.BsrD1 nicking sites. Finally, to generate p3d-Phen-A, the vector was nicked using Nt. BspQ1 and Nb.BsrD1 and ligated with an excess of the following oligo containing a 3-deaza-3-phenethyl-adenosine (3d-Phen-A) at the alanine on position 15: 5′-ACCTCAACTACTTG**A**CCCTCCTCATT-3’ (Malvezzi et al., 2017). pAP-ICL was generated as previously described (Semlow et al., 2016, 2022).

### Antibodies and Immunodepletions

Antibodies against Rev1 (Rev1-N and Rev1-C) (Budzowska et al., 2015), Rfwd3 (Gallina et al., 2021), Polη (Gallina et al., 2021), and HMCES (Semlow et al., 2022) were described previously. Polκ, Rev3, and FancA antibodies were raised by New England Peptide by immunizing rabbits with Ac-CPASKKSKPNSSKNTIDRFFK-OH, Ac-CLADLSIPQLD-OH, and Ac-CSFKAPDDYDDLFFEPVF-OH, respectively.

To immunodeplete Rev1 from *Xenopus* egg extracts, an equal of Protein A Sepharose Fast Flow (PAS) (GE Health Care) beads was bound to Rev-N or Rev1-C antibodies overnight at 4°C. The beads were then washed twice with 500 µL PBS, once with ELB (10 mM HEPES pH 7.7, 50 mM KCl, 2.5 mM MgCl2, and 250 mM sucrose), twice with ELB supplemented with 0.5 M NaCl, and twice with ELB. One volume of precleared HSS or NPE was then depleted by mixing with 0.2 volumes of antibody-bound beads and then incubated at room temperature for 15 min, before being harvested. For HSS, the depletion procedure was performed once with Rev1-N coupled beads and once with Rev1-C coupled beads. For NPE the depletion procedure was performed twice with Rev1-N coupled beads and once with Rev1-C coupled beads. To immunodeplete Polκ, Polη or RFWD3 from *Xenopus* egg extracts, one volume of PAS beads was bound to five volumes of affinity purified antibody (1 mg/mL). The beads were washed as described above, and one volume of precleared HSS or NPE was then depleted by mixing with 0.2 volumes of antibody-bound beads for 15 min at room temperature. The depletion procedure was performed once for HSS and three times for NPE. For HMCES and Polκ combined depletion, one volume of beads was bound to 8 volumes of each affinity purified antibody (1 mg/mL). The beads were washed and depletion was performed as described for Polκ immunodepletion.

### Immunoprecipitations

For the FancA and Polκ immunoprecipitation experiments, 5 μL of PAS beads were incubated with 10 μg of the respective affinity purified antibody for 1h at RT. The sepharose beads were subsequently washed twice with PBS and three times with IP buffer 1 (10 mM Hepes pH 7.7, 50 mM KCl, 2.5 mM MgCl2, 0.25% NP-40). Next, 5 μL of NPE was diluted with 20 μL of IP buffer and incubated with antibody prebound beads for 1 hour at RT. The beads were then washed three times with IP buffer and resuspended in 50 μL of 2x Laemmli sample buffer before analysis by Western blotting.

### Nascent leading strand analysis

For nascent leading strand analysis, 3-4 µL of replication reaction were added to 10 volumes of transparent stop buffer (50 mM Tris-HCl, pH 7.5, 0.5% SDS, 25 mM EDTA). The replication intermediates were purified as previously described (Knipscheer et al., 2009; Räschle et al., 2008). The DNA was digested with the indicated restriction enzymes and subsequently supplemented with 0.5 volumes of denaturing PAGE Gel Loading Buffer II (Life technologies). The digested DNA products were separated on a 6% polyacrylamide sequencing gel.

### Protein expression and purification

Full length *Xenopus laevis* Polκ with an N-terminal 6xHis-tag was amplified from pCMV-Sport.ccdb-Polκ (Williams et al., 2012) and cloned into pET28b (Novagen) using primers A and B, and restriction enzymes BamHI and XhoI. *Xenopus* Polκ C-ter with an N-terminal 6xHis-tag was cloned into pET28b using primers B and C, and restriction enzymes BamHI and XhoI. Polκ mutations were introduced via Quikchange mutagenesis and confirmed by Sanger sequencing.

Plasmids containing either Polκ wild-type, Polκ mutants, or Polκ C-ter were transformed into BL21 *E.coli* competent cells. Cells were grown at 37°C to O.D. 0.6-0.8 in LB broth and were subsequently inducted with 0.5 mM IPTG for 4h. Bacteria were harvested by centrifugation and resuspended in 20 mL of lysis buffer (50 mM Tris pH 7.5, 300 mM NaCl, 2 mM MgCl_2_ and 1 mM DTT, 1X Roche EDTA-free Complete protease inhibitor cocktail). Suspensions were sonicated and cleared by high-speed centrifugation 15000 rpm in a F15-8x50cy rotor for 1hr at 4°C. The soluble fraction was collected and incubated with 2 mL of Ni-NTA Superflow affinity resin (Qiagen), previously equilibrated with lysis buffer, for 2h at 4°C. The resin was then washed three times with 20 mL of Wash Buffer (50 mM Tris pH 7.5, 300 mM NaCl, 2 mM MgCl_2_, 1 mM DTT, 0,1% Triton-X, 10 mM imidazole). 6xHis-Polκ was subsequently eluted with Elution Buffer (50 mM Tris pH 7.5, 300 mM NaCl, 2 mM MgCl_2_, 1 mM DTT, 10% glycerol, 10 mM imidazole). Elution fractions containing the target proteins were pooled and dialyzed against dialysis buffer (50 mM Tris pH 7.5, 300 mM NaCl, 2 mM MgCl_2_, 1 mM DTT, 10% glycerol) at 4°C overnight. After dialysis, protein fractions were concentrated to 100 μL using 30,000 MWCO centrifugal filters (Amicon) and subsequently aliquoted, flash-frozen in liquid nitrogen and stored at − 80°C.

Primer A: 5’ – ATGCGGATCCAATGGATAACAAGCAAGAAGCAGAG – 3’

Primer B: 5’ – ATGCCTCGAGCTACTTGAAGAATCTGTCGATGGTG – 3’

Primer C: 5’ – ATGCGGATCCAAAACATCACCAGAAGAGCATTACTAG – 3’

Plasmids for expressing *Xenopus laevis* Polκ wild-type (WT) and catalytic dead (CD) in rabbit reticulocytes were kind gifts from Prof. Jean Gautier (Williams et al., 2012). Briefly, 2 μg of pCMV-Sport- Polκ were incubated with 100 µL of TnT® Sp6 Quick Master Mix (Promega) supplemented with 4 μL of 1 mM methionine for 90 min at 30°C. The reaction volume was subsequently adjusted to 400 μL with PBS and DNA was precipitated by addition of 0.06% polymin-P and incubation for 30 min at 4°C with rotation. The mixture was then centrifuged at 14000 g for 30 min and the proteins in the supernatant were precipitated with saturated ammonium sulfate to a final concentration of 55% for 30 min at 4°C with rotation, followed by centrifugation at 16000 g for 30 min. The protein pellet was subsequently resuspended in 15 μL of ELB buffer, dialyzed for 3 hours at 4°C in ELB buffer. As a negative control, a reaction without DNA was performed. The Polκ protein preparations obtained by this method was used for gap filling synthesis experiments (Figure S1D).

### Chromatin spin-down

Demembranated *Xenopus* sperm chromatin was prepared as described (Sparks & Walter, 2019) and stored at −80°C at a concentration of 100000 sperm chromatin/µL (320 ng/µL). For analysis of UV-damaged chromatin, sperm chromatin was diluted to either 25000 sperm chromatin/µL in ELB buffer (10 mM HEPES pH 7.7, 50 mM KCl, 2.5 mM MgCl_2_, and 250 mM sucrose), deposited on parafilm and irradiated with 2000 J/m^2^ of UV-C. For non-replicating reactions, HSS and NPE were mixed at a 1:2 ratio. Subsequently, undamaged or UV-damaged sperm chromatin was added at a final concentration of 16 ng/ µL to HSS/NPE mix that was previously mock- Polη-, Polκ-, Rev1 or Rfwd3-depleted . At the indicated timepoints, 8 µL of replication reaction was stopped with 60 µL of ELB buffer supplemented with 0.2% Triton-X. The mixture was carefully layered on top on a sucrose cushion (10 mM HEPES pH 7.7, 50 mM KCl, 2.5 mM MgCl2, and 500 mM sucrose) and spun for 1 min at 6800 x g in a swing bucket centrifuge at 4°C. The chromatin pellet was carefully washed twice with 200 µL of ice-cold ELB buffer and resuspended in 2X Laemmli buffer.

### Alphafold models generation

Molecular models were predicted using AlphaPulldown 2.3.1 (Yu et al., 2023), running AlphaFold 0.30.0 (Jumper et al., 2021). AlphaPulldown parameters were as follows: cycles=3, models=5, predictions/model=1. Structure predictions were generated for *Xenopus laevis* Q6DFE4 (POLK), P18248 (PCNA), Q6NRK6 (REV1), D0VEW8 (REV3), Q8QFR4 (REV7), O93610 (POLD2), Q76LD3 (POLD3), and P62972 (UBIQP), individually and as complexes of either full-length proteins or protein fragments.

Models were evaluated on their Predicted Local Distance Difference Test (Jumper et al., 2021), Interface Predicted Template Modelling (Evans et al., 2022; Yu et al., 2023), Predicted Template Modeling (Jumper et al., 2021) and Predicted Aligned Error (Varadi et al., 2022) scores. From each prediction, the best model as determined by Alphapulldown, was selected for inclusion in the final complex models.

Model building was performed using UCSF ChimeraX (Goddard et al., 2018; Pettersen et al., 2021). The catalytic complex was modelled on a scaffold of Human Pol Kappa holoenzyme with Ub-PCNA (**PDB: 7NV1**, Lancey et al., 2021).

The non-catalytic complex was modelled on a scaffold of the yeast Polζ (**PDB: 6V93**, Malik et al., 2020). To establish the relative position of Polζ to PCNA a structure of Processive human polymerase delta holoenzyme was used (PDB: **6TNY**, Lancey et al., 2020). The monoubiquitinated PCNA and scaffold DNA attached to the polymerase complex was modelled on a structure of Mono-ubiquitinated PCNA (PDB: **3TBL**, Zhang et al., 2012)

### CHROMASS

CHROMASS experiments were performed as previously described (Räschle et al., 2015). Briefly, isolated sperm chromatin was either untreated or treated with 2000 J/m^2^ of UV-C. Each reaction was performed in quadruplicate. The sperm chromatin was then incubated at a final concentration of 16 ng/µL in non-licensing extracts that were either mock-, Polκ- or Rev1-depleted. Reactions were stopped after 45 min. Specifically, 10 µL of replication reaction was stopped with 60 µL of ELB buffer supplemented with 0.2% Triton-X, and chromatin spin down performed as described above. The chromatin pellet was then resuspended in 100 µl denaturation buffer (9 M urea, 100 mM Tris-HCl, pH 8), and transferred to a new low binding tube. Cysteines were reduced (1 mM DTT for 15 min at RT) and alkylated (0.55 M chloroacetamide for 40 min at RT protected from light). Proteins were first digested with 0.5 µg LysC (2.5 hours at RT) and then with 0.5 µg tripsin at 30°C overnight. Peptides were acidified with 10% trifluoroacetic acid (pH <4), followed by addition of 400 mM NaCl, and purified by StageTip (C18 material). For this, StageTips were first activated in 100% methanol, then equilibrated in 80% acetonitrile in 0.1% formic acid, and finally washed twice in 0.1% formic acid. Samples were loaded on the equilibrated stage tips and washed twice with 50 µL 0.1% formic acid. StageTip elution was performed with 80 μL of 25% acetonitrile in 0.1% formic acid, eluted samples were dried to completion in a SpeedVac at 60 °C, dissolved in 10 μL 0.1% formic acid, and stored at −20°C until MS analysis.

### MS data acquisition

All MS samples were analyzed on an EASY-nLC 1200 system (Thermo) coupled to an Orbitrap Exploris™ 480 mass spectrometer (Thermo). Out of the n=4 biochemical replicates, 50% was analyzed per run (R1-R4). Afterwards, an additional set of n=4 technical replicates was performed by mixing 25%:25% of R1:R2 (=R5), R2:R3 (=R6), R3:R4 (=R7), and R4:R1 (=R8), totaling n=8 technical replicates. Separation of peptides was performed using 20-cm columns (75 μm internal diameter) packed in-house with ReproSil-Pur 120 C18-AQ 1.9 μm beads (Dr. Maisch). Elution of peptides from the column was achieved using a gradient ranging from buffer A (0.1 % formic acid) to buffer B (80 % acetonitrile in 0.1 % formic acid), at a flow of 250 nl/min. The gradient length was 80 min per sample, including ramp-up and wash-out, with an analytical gradient of 58 min ranging from 7% B to 34% B. Analytical columns were heated to 40°C using a column oven, and ionization was achieved using a NanoSpray Flex™ NG ion source. Spray voltage set to 2 kV, ion transfer tube temperature to 275°C, and RF funnel level to 40 %. Full scan range was set to 300-1,300 m/z, MS1 resolution to 120,000, MS1 AGC target to “200” (2,000,000 charges), and MS1 maximum injection time to “Auto”. Precursors with charges 2-6 were selected for fragmentation using an isolation width of 1.3 m/z, and fragmented using higher-energy collision disassociation (HCD) with normalized collision energy of 25. Precursors were excluded from re-sequencing by setting a dynamic exclusion of 80 s. MS2 AGC target was set to “200” (200,000 charges), intensity threshold to 360,000 charges per second, MS2 maximum injection time to “Auto”, MS2 resolution to 30,000, and number of dependent scans (TopN) to 13.

### MS data analysis

All MS RAW data were analyzed using the freely available MaxQuant software (Cox & Mann, 2008), v. 1.5.3.30, in a single computational run. Default MaxQuant settings were used, with exceptions specified below. For generation of theoretical spectral libraries, the *Xenopus laevis* FASTA database was downloaded from UniProt on the 3rd of October 2022. *In silico* digestion of proteins to generate theoretical peptides was performed with trypsin, allowing up to 3 missed cleavages. Minimum peptide length was set to 6, and maximum peptide mass was set to 6,000 Da. Allowed variable modifications were oxidation of methionine (default), protein N-terminal acetylation (default), deamidation of asparagine and glutamine, peptide N-terminal glutamine to pyro-glutamate conversion, di-oxidation of tryptophan, and replacement of three protons by iron (cation Fe[III]) on aspartate and glutamate. These variable modifications were determined by an initial analysis of the RAW data using pFind v3.1.6 in “Open search” mode (Chi et al., 2018), to unbiasedly determine any known modifications (from the Unimod database) affecting >0.5% of peptide-spectrum matches (PSMs) across all samples. Maximum variable modifications per peptide were set to 3. Label-free quantification (LFQ) using MaxLFQ was enabled (Cox et al., 2014) with “Fast LFQ” disabled. Matching between runs was enabled, with an alignment window of 20 min and a match time window of 1 min. Stringent MaxQuant 1% FDR control was applied at the PSM-, protein-, and site-decoy levels (default).

### MS data annotation and quantification

The *Xenopus laevis* FASTA databases downloaded from UniProt lacked comprehensive gene name annotations. Missing or uninformative gene names were, when possible, semi-automatically curated, as described previously (Gallina et al., 2021). Quantification of the MaxQuant output files (“proteinGroups.txt”), and all statistical handling, was performed using Perseus software v1.5.5.3 (Tyanova et al, 2016). In total, n=8 technical replicates (derived from n=4 biochemical replicates) were analyzed. For quantification purposes, all LFQ-normalized protein intensity values were log2 transformed, and filtered for presence in 8 of 8 replicates (n = 8/8) in at least one experimental condition. Missing values were imputed below the global experimental detection limit at a downshift of 1.8 and a randomized width of 0.15 (in log2 space). Statistical significance of differences was in all cases tested using two-tailed Student’s two-sample t-testing, with permutation-based FDR control applied to ensure a corrected p-value (i.e. q-value) of <1%. Proteins not enriched over the no-DNA control in at least one CHROMASS condition (FDR<1%, s0=1, 2,500 rounds of randomization) were removed from the analysis, after which previously imputed values were re-imputed based on the new total matrix. Final biological differences were determined using two-tailed Student’s two-sample t-testing (FDR<1%, s0=0.5, 2,500 rounds of randomization) on the remaining CHROMASS-enriched proteins.

The mass spectrometry proteomics data have been deposited to the ProteomeXchange Consortium via the PRIDE (Perez-Riverol et al., 2022) partner repository with the dataset identifier PXD044258.

Reviewer account details:

Username

Password: vkOh8JQ3

### Base editor tiling screen and analysis

A single guide RNA (sgRNA) library predicted to introduce missense mutations in the coding sequence of POLK was designed, synthesized (GenScript), and cloned into the Abe8e-SpG lentiviral vector pRDA_479. pRDA_479 was a gift from John Doench & David Root (Addgene plasmid #179099, Sangree et al., 2022). The POLK tiling library was part of a larger tiling library for which only POLK is analyzed here. Lentiviral particles were produced by co-transfection of the sgRNA library with lentiviral packaging plasmids pMD2G and psPAX2 in HEK293T/17 cells using Lipofectamine 3000 (Invitrogen). pMD2.G and psPAX2 were gifts from Didier Trono (Addgene plasmid #12259 and #12260). 6 hours after transfection, medium was exchanged for DMEM GlutaMax + 10% FBS + 100 U/mL penicillin–streptomycin (Gibco) + 1% bovine serum albumin. 48 hours after transfection, viral particles were harvested and filtered through a 0.45 μm syringe filter before freezing at −80°C. RPE1-hTERT p53-/- cells (a kind gift from D. Durocher) were cultured in DMEM GlutaMax supplemented with 10% FBS and 100 U/mL penicillin–streptomycin (Gibco) and passaged every three days. The screen was performed as a duplicate (two separate transductions). Cells were transduced with the lentiviral library at a low multiplicity of infection (0.3-0.4) at a coverage of >500-fold sgRNA representation, which was maintained throughout the screen. Transductions were performed by treating cells with 8μg/mL polybrene and lentiviral supernatants for 24 hours. Transduced cells were selected by treatment with 20 μg/mL puromycin for 24 hours followed by trypsinization and reseeding in the same plates with 20 μg/mL puromycin for another 24 hours. After selection, cells were passaged for 6 days before splitting into untreated or Illudin S treated fractions where they were passaged for an additional 12 days with passaging every 3 days in medium with or without a low dose of Illudin S (1.4 ng/mL) equivalent to predetermined LD_20_ concentrations in uninfected cells (actual LDs obtained during the screen were LD_13_ and LD_7_ for replicate 1 and 2). Genomic DNA was extracted from cell pellets harvested 3 days after transduction (t0) and at the final timepoint (t18). The genomic DNA region containing the integrated sgRNA was amplified by PCR using Q5 Mastermix Next Ultra II (New England Biolabs) with the LCV2_forward and LCV2_reverse primers (**Table S3**). A second PCR reaction introduced i5 and i7 multiplexing barcodes (**Table S3**) and gel-purified products were sequenced on Illumina NextSeq2000. Sequencing data of t0, untreated t18, Illudin S treated t18 samples was converted to gRNA sequencing counts by mapping to sgRNAs tiling POLK. Two replicates were collected per condition resulting in two data points per guide per condition. Low abundance sgRNAs were removed (counts < 30) and raw sequencing counts were normalized per condition replicate to log2 transcripts per million (log2TPM). The log2TPM values were compared using limma (Ritchie et al., 2015) for three sample pairs: t0 vs untreated t18; t0 vs Illudin S treated t18; and untreated t18 vs Illudin S treated t18 and the fold change, p-value and corrected p-value were collected for each gRNA.

## Author Contributions

S.S.B. performed all experiments unless stated otherwise. A.C. generated the AlphaFold models in Figures 1C, S1B-C, 7 and S7 under the supervision of T.M. S.M.A. and B.M. performed the CRISPR base editor tiling screen (Figure 1B, Figure S1A) under the supervision of J.N. E.P.T.H. provided technical advice on base editor screen. N.D. performed statistical analysis of the sequencing data. I.A.H. performed MS analysis on CHROMASS samples (Figure 6 and S6), under the supervision of M.L.N. L.S and B.B. generated the p3d-Phen-A minor groove DNA substrate under the supervision of S.S and J.P.D. I.G. performed the experiment in Figure 6D and preliminary experiments for this project. D.S. provided the AP-ICL containing plasmid. S.S.B. and J.P.D. designed and analyzed the experiments performed in *Xenopus* egg extracts. S.S.B. and J.P.D. prepared the manuscript with feedback and input from all authors of the manuscript.

## Acknowledgments

We thank Jean Gautier for sharing Polκ expressing constructs, Daniel Durocher for the RPE1-hTERT p53-/- cells, and staff of the CPR/ReNew Genomics Platform for support: H. Wollmann, M.Michaut, A. Kalvisa. We also thank members of the Duxin laboratory for feedback on the manuscript. The Novo Nordisk Foundation Center for Protein Research is supported financially by the Novo Nordisk Foundation (grant agreement NNF14CC0001). This project has received funding from the European Research Council (ERC) under the European Union’s Horizon 2020 research and innovation program (grant agreement 715975) and from the Novo Nordisk Foundation (grant NNF22OC0074140). T.C.R.M. was supported by a Novo Nordisk Fonden Hallas-Møller Emerging Investigator Grant (NNF22OC0073571), the Danish National Research Foundation (DNRF115), and the Carlsberg Foundation (CF21-0571).

## Declaration of Interests

The authors declare no competing interests.

## Supplemental Figure Legends

**Figure S1. A**) Dot plot showing the results of the Polκ CRISPR base editor tiling screen comparing t0 and t18 of the untreated condition. Each guide is shown as a dot. X-axis represents amino acid position in Polκ. Y-axis represents Log2 fold changes between t0 and t18 of the untreated condition. Larger dots represent guides which are significantly changing between the two conditions (p-value ≤ 0.01). The various domains of Polκ are indicated. **B**) Composite molecular model of human Polκ. Dashed lines represent disordered regions that are not present in the model. The model was generated as described in Figure 1C. **C**) Composite molecular model of human Polκ with point mutations derived from the base editor tiling screen. Dashed lines represent disordered regions that are not present in the model. Point mutations are highlighted in red. The left panel is a duplicate of Figure 1C. **D**) pCTRL and p3d-PhenA were replicated in the presence or absence of ubiquitin E1 inhibitor. Reaction samples were analyzed as in Figure 1G. **E**) Schematic diagram illustrating the replication intermediates generated during the replication of p3d-PhenA. **F**) Schematic representation of gap filling synthesis when pDPC^ssDNA^ is incubated with Polκ WT, Polκ CD, or non-licensing egg extracts (non-replicative). Note the absence of DNA synthesis in the Polκ CD reaction.

**Figure S2. A**) Polκ-depleted extracts were compared to a mock depletion dilution series. Polκ-depleted extracts were supplemented with either buffer (+Buffer), wild-type Polκ (+WT), or RIR (RIR*) Polκ mutant. Samples were blotted with a Polκ antibody. **B**) Extracts from (A) were used to replicate p3d-Phen-A. Samples were analyzed as in Figure 1G. **C**) Polκ-depleted extracts were supplemented with either buffer (+Buffer), wild-type Polκ (+WT), PIP1 (PIP1*), or PIP2 (PIP2*) Polκ mutants. Samples were blotted with Polκ and 6x-His antibodies. Note that the Polκ antibody was generated against Polκ C-terminal end which encompasses PIP2. This antibody exhibits lower affinity for Polκ PIP2* compared to Polκ WT and Polκ PIP1*. **D**) Extracts from (C) were used to replicate p3d-Phen-A. Samples were analyzed as in Figure 1G. **E**) Polκ-depleted extracts were supplemented with either buffer (+Buffer), wild-type Polκ (+WT), UBZ (UBZ1*), or UBZ (UBZ2*) Polκ mutants. Samples were blotted with a Polκ antibody. **F**) Extracts from (E) were used to replicate 3d-Phen-A. Samples were analyzed as in Figure 1G.

**Figure S3. A**) Replication intermediates generated during replication of pDPC (Duxin et al., 2014). **B)** Mock-, Polη-, Polκ-, Rev1-, or Rev3-depleted extracts were blotted with the indicated antibodies. **C)** pDPC was replicated in egg extracts in mock-, Polη-, Polκ-, or Rev1-depleted extracts. Reaction samples were analyzed as in Figure 1G. **D)** pDPC was replicated in Polκ- or Rev1-, or Rev3-depleted egg extracts. Samples were digested, separated on a denaturing polyacrylamide gel and analyzed as in Figure 3C.

**Figure S4. A)** Generation of pDPC^PK^ (Larsen et al., 2019). **B)** Mock- and Polκ- depleted egg extracts were used to replicate pDPC^PK^. Polκ-depleted extracts were supplemented with either buffer (+Buffer), wild-type (+WT), or catalytically inactive (+CD) Polκ. Samples were analyzed as in Figure 1G. **C)** Samples from (B) were digested and analyzed as in Figure 3C. **D)** pDPC^Lead^ was replicated in either mock- or Polκ-depleted egg extracts in the presence of LacI (Duxin et al., 2014). Polκ-depleted extracts were supplemented with either buffer (+Buffer), wild-type Polκ (+WT), or catalytically inactive Polκ (+CD). Samples were digested with AatII, separated on a denaturing polyacrylamide gel and analyzed as in Figure 3C. **E)** Extracts from (D) were used to replicate pDPC^Lag^ (Duxin et al., 2014). Samples were digested with BssHII, separated on a denaturing polyacrylamide gel and analyzed as in Figure 3C. **F**) Mock- and Polκ-depleted egg extracts were supplemented with either buffer (+Buffer), recombinant Polκ wild-type (+WT), PIP1 (+PIP1*), or PIP2 (+PIP2) Polκ mutants. Extracts were then used to replicate pDPC. Samples were digested and analyzed as in Figure 3C. **G**) Mock- and Polκ-depleted egg extracts were supplemented with either buffer (+Buffer), UBZ1(+UBZ1*), UBZ2 (+UBZ2*), or UBZ1 and UBZ2 (+UBZ1+2*) Polκ mutants. Extracts were then used to replicate pDPC. Samples were analyzed as in Figure 1G.

**Figure S5**. **A)** Mock- and Polκ-depleted egg extracts were used to replicate pAP-ICL. Polκ-depleted extracts were supplemented with either buffer (+Buffer), wild-type Polκ (+WT), or catalytically inactive Polκ (+CD). The samples were analyzed as in Figure 1G. **B)** Mock- and HMCES-extracts were blotted with a HMCES antibody. * indicates a non-specific band. **C)** HMCES-depleted extracts were either mock- or Polκ-depleted and were subsequently used to replicate pAP-ICL. Polκ-depleted extracts were supplemented with either buffer (+Buffer), wild-type Polκ (+WT), or catalytically inactive Polκ (+CD). The samples were analyzed as in Figure 1G.

**Figure S6. A)** Mock-, Polη-, or double Polη- and Polκ-depleted egg extracts were used to replicate pDPC. Polκ-depleted extracts were supplemented with either buffer (+Buffer), wild-type Polκ (+WT), or catalytically inactive Polκ (+CD). Samples were analyzed as in Figure 1G. **B)** MS analysis of protein recruitment to UV-treated sperm chromatin compared to untreated sperm chromatin in egg extracts. The volcano plot shows the difference in abundance of proteins between the two sample conditions (x-axis), plotted against the *p*-value resulting from two-tailed Student’s two-sample t-testing (y-axis). Proteins significantly down- or up-regulated (FDR<5%) upon UV treatment are represented in red or blue, respectively. TLS polymerases are highlighted in orange. n=4 biochemical and n=8 technical replicates, significance was determined via two-tailed Student’s two-sample t-testing, with permutation-based FDR control (s0=0.5) to ensure an adjusted p-value (i.e. q-value) of <0.05 in all cases. The significance line is drawn at q=0.01. **C)** Same experiment as in (B) but comparing mock- to Rev1-depleted extracts. Small red dots, 1% < FDR < 5%; large red dots, FDR < 1%. Polκ and Rev1 are highlighted in purple and black, respectively. Note that only proteins that were significantly enriched in (B) are visualized. **D)** Same experiment as in (B) but comparing mock- to Polκ-depleted extracts. Small red dots, 1% < FDR < 5%; large red dots, FDR < 1%. Polκ and Rev1 are highlighted in purple and black, respectively. Note that only proteins that were significantly enriched in (B) are visualized. **E)** Western blot of the co-immunoprecipitation of Polκ with FancA. SUP, supernatant; IP, immunoprecipitation. Note that the FA pathway is not involved in the repair nor bypass of M.HpaII DPCs (Duxin et al., 2014).

**Figure S7. A)** AlphaPulldown predictions of protein complexes were utilised to construct the composite molecular models presented in Figure 7. Each panel is divided into three sections: The upper panel displays the Predicted Aligned Error (PAE) plots associated with specific protein/fragment combinations generated by AlphaPulldown (Yu et al., 2023). The PAE score in AlphaFold estimates the distance errors for residue pairs. It is represented by plots composed by diagonal squares for inner- correlation and cross-correlation areas for protein/fragment interactions. They assess prediction confidence, with low values being reliable and high values unreliable. The middle panel shows the predicted structures. Lastly, the lower panel includes the names of the predicted protein combinations, along with their corresponding Interface Predicted Template Modelling (IPTM), the Predicted Template Modelling (PTM) scores and the Ranking of the model (R0). **B)** Summary tables featuring the protein structures used as scaffolds for model building, along with their respective PDB codes. Single protein predictions obtained from AlphaPulldown with related Predicted Local Distance Difference Test scores (pLDDT). Uniprot accession codes for each of the proteins used for the complex protein modelling.

## Supplemental Table Legends

**Table S1. POLK base editor tiling screen results and analysis.** Related to Figure 1 and S1.

**Table S2. MS analysis of protein recruitment to UV-treated sperm chromatin in mock-, Rev1-, or Pol**κ**-depleted extracts.** Related to Figure 6 and S6. ΔMock, mock-depleted extracts incubated in undamaged chromatin; ΔMock +UV, mock-depleted extracts incubated with UV-treated chromatin; ΔRev1 +UV, Rev1-depleted extracts incubated with UV-treated chromatin; ΔPolκ +UV, Polκ-depleted extracts incubated with UV-treated chromatin.

**Table S3. PCR primers used during base editor tiling screen**. Related to Figure 1 and S1 (see Materials and methods).

## Notes

### Competing Interest Statement

The authors have declared no competing interest.

