## Supplementary material for "Catalytic and non-catalytic functions of DNA polymerase kappa in translesion DNA synthesis": Sup Figures

Figure S1

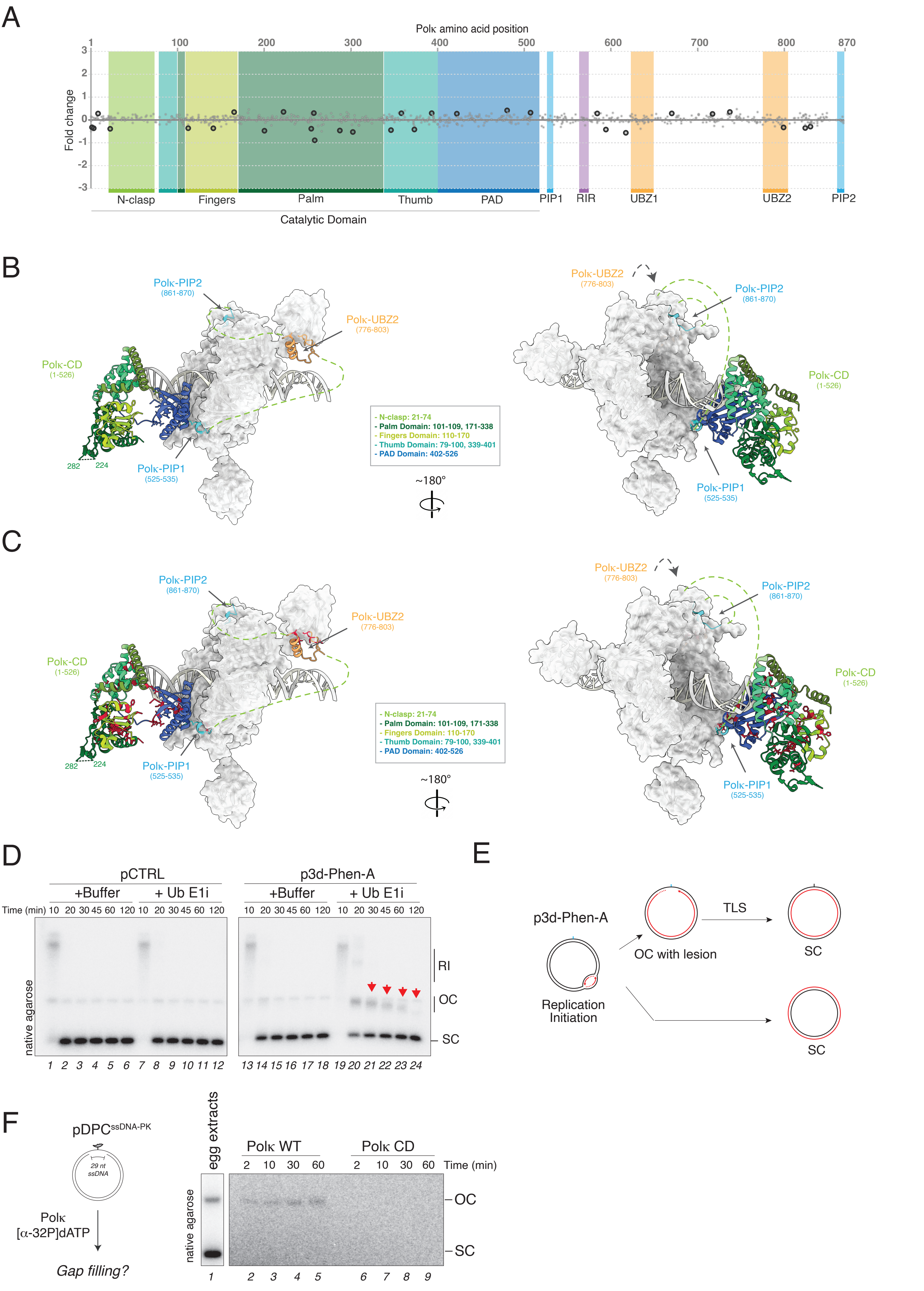
A
Polκ amino acid position
1
100
200
300
400
500
600
700
800
870
Fold change
N-clasp
Fingers
Palm
Thumb
PAD
PIP1
RIR
UBZ1
UBZ2
PIP2
Catalytic Domain
B
Polκ-PIP2 (861-870)
Polκ-UBZ2 (776-803)
Polκ-CD (1-526)
Polκ-PIP1 (525-535)
282
224
- N-clasp: 21-74
- Palm Domain: 101-109, 171-338
- Fingers Domain: 110-170
- Thumb Domain: 79-100, 339-401
- PAD Domain: 402-526
~180°
C
D
pCTRL
p3d-Phen-A
+Buffer
+ Ub E1i
Time (min)
native agarose
RI
OC
SC
E
p3d-Phen-A
Replication Initiation
OC with lesion
TLS
SC
F
pDPC ssDNA-PK
29 nt ssDNA
Polκ
[α-32P]dATP
Gap filling?
egg extracts
Polκ WT
Polκ CD
Time (min)
OC
SC

Figure S2

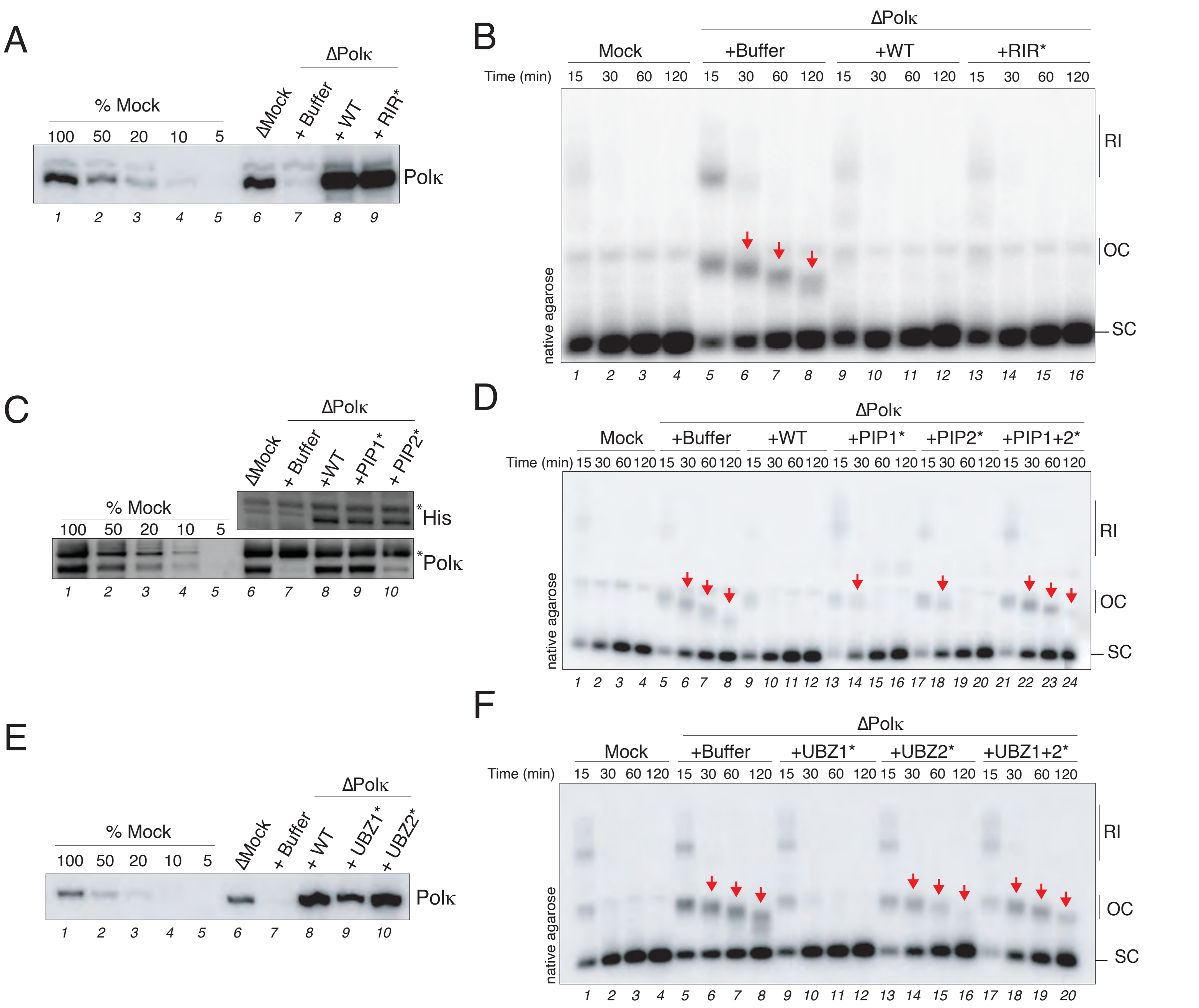

A
% Mock
100 50 20 10 5
ΔPolκ
ΔMock
+ Buffer
+ WT
+ RIR*
Polκ
1 2 3 4 5 6 7 8 9
B
ΔPolκ
Mock
+Buffer
+WT
+RIR*
Time (min) 15 30 60 120 15 30 60 120 15 30 60 120 15 30 60 120
native agarose
RI
OC
SC
1 2 3 4 5 6 7 8 9 10 11 12 13 14 15 16
C
ΔPolκ
ΔMock
+ Buffer
+WT
+PIP1*
+ PIP2*
% Mock
100 50 20 10 5
*His
*Polκ
1 2 3 4 5 6 7 8 9 10
D
ΔPolκ
Mock
+Buffer
+WT
+PIP1*
+PIP2*
+PIP1+2*
Time (min) 15 30 60 120 15 30 60 120 15 30 60 120 15 30 60 120 15 30 60 120 15 30 60 120
native agarose
RI
OC
SC
1 2 3 4 5 6 7 8 9 10 11 12 13 14 15 16 17 18 19 20 21 22 23 24
E
ΔPolκ
% Mock
100 50 20 10 5
ΔMock
+ Buffer
+ WT
+ UBZ1*
+ UBZ2*
Polκ
1 2 3 4 5 6 7 8 9 10
F
ΔPolκ
Mock
+Buffer
+UBZ1*
+UBZ2*
+UBZ1+2*
Time (min) 15 30 60 120 15 30 60 120 15 30 60 120 15 30 60 120 15 30 60 120
native agarose
RI
OC
SC
1 2 3 4 5 6 7 8 9 10 11 12 13 14 15 16 17 18 19 20

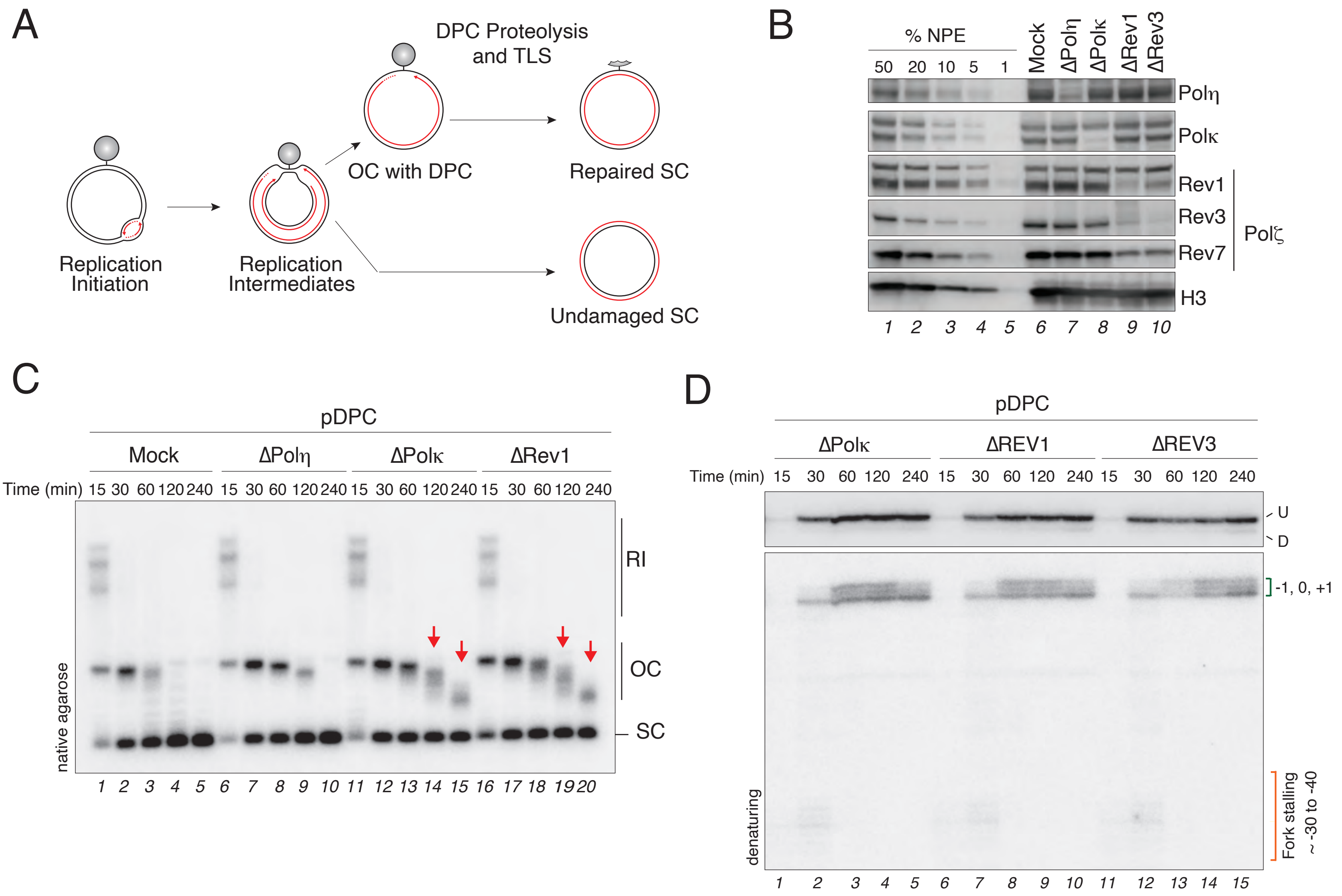
A
DPC Proteolysis and TLS
OC with DPC
Repaired SC
Replication Initiation
Replication Intermediates
Undamaged SC
B
% NPE
50 20 10 5 1
Mock ΔPolη ΔPolκ ΔRev1 ΔRev3
Polη
Polκ
Rev1
Rev3
Rev7
Polζ
H3
1 2 3 4 5 6 7 8 9 10
C
pDPC
Mock ΔPolη ΔPolκ ΔRev1
Time (min) 15 30 60 120 240 15 30 60 120 240 15 30 60 120 240 15 30 60 120 240
RI
OC
SC
native agarose
1 2 3 4 5 6 7 8 9 10 11 12 13 14 15 16 17 18 19 20
D
pDPC
ΔPolκ ΔREV1 ΔREV3
Time (min) 15 30 60 120 240 15 30 60 120 240 15 30 60 120 240
U
D
-1, 0, +1
Fork stalling
~ -30 to -40
denaturing
1 2 3 4 5 6 7 8 9 10 11 12 13 14 15

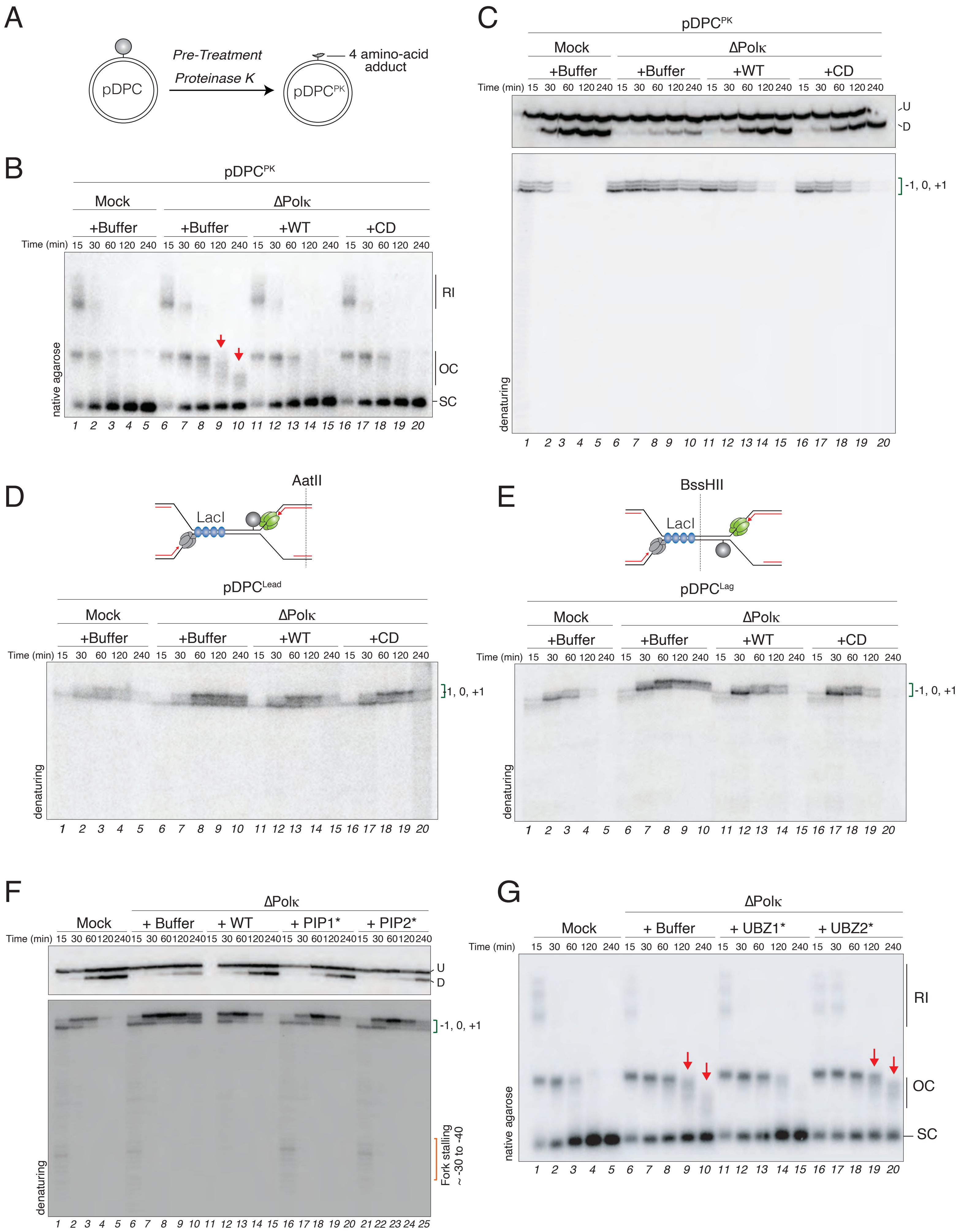

A
pDPC
Pre-Treatment
Proteinase K
pDPC^PK
4 amino-acid adduct
B
pDPC^PK
Mock
ΔPolκ
+Buffer
+Buffer
+WT
+CD
Time (min) 15 30 60 120 240 15 30 60 120 240 15 30 60 120 240 15 30 60 120 240
RI
OC
SC
native agarose
1 2 3 4 5 6 7 8 9 10 11 12 13 14 15 16 17 18 19 20
C
pDPC^PK
Mock
ΔPolκ
+Buffer
+Buffer
+WT
+CD
Time (min) 15 30 60 120 240 15 30 60 120 240 15 30 60 120 240 15 30 60 120 240
U
D
-1, 0, +1
denaturing
1 2 3 4 5 6 7 8 9 10 11 12 13 14 15 16 17 18 19 20
D
AatII
LacI
pDPC^Lead
Mock
ΔPolκ
+Buffer
+Buffer
+WT
+CD
Time (min) 15 30 60 120 240 15 30 60 120 240 15 30 60 120 240 15 30 60 120 240
1, 0, +1
denaturing
1 2 3 4 5 6 7 8 9 10 11 12 13 14 15 16 17 18 19 20
E
BssHII
LacI
pDPC^Lag
Mock
ΔPolκ
+Buffer
+Buffer
+WT
+CD
Time (min) 15 30 60 120 240 15 30 60 120 240 15 30 60 120 240 15 30 60 120 240
-1, 0, +1
denaturing
1 2 3 4 5 6 7 8 9 10 11 12 13 14 15 16 17 18 19 20
F
ΔPolκ
Mock
+ Buffer
+ WT
+ PIP1*
+ PIP2*
Time (min) 15 30 60 120 240 15 30 60 120 240 15 30 60 120 240 15 30 60 120 240 15 30 60 120 240
U
D
-1, 0, +1
Fork stalling
~ -30 to -40
denaturing
1 2 3 4 5 6 7 8 9 10 11 12 13 14 15 16 17 18 19 20 21 22 23 24 25
G
ΔPolκ
Mock
+ Buffer
+ UBZ1*
+ UBZ2*
Time (min)
15 30 60 120 240 15 30 60 120 240 15 30 60 120 240 15 30 60 120 240
RI
OC
SC
native agarose
1 2 3 4 5 6 7 8 9 10 11 12 13 14 15 16 17 18 19 20

Figure S5

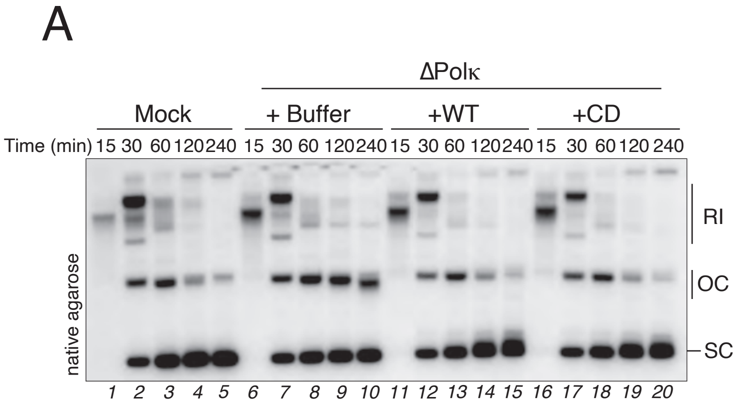

A
ΔPolκ
Mock
+ Buffer
+WT
+CD
Time (min) 15 30 60 120 240 15 30 60 120 240 15 30 60 120 240 15 30 60 120 240
native agarose
RI
OC
SC
1 2 3 4 5 6 7 8 9 10 11 12 13 14 15 16 17 18 19 20


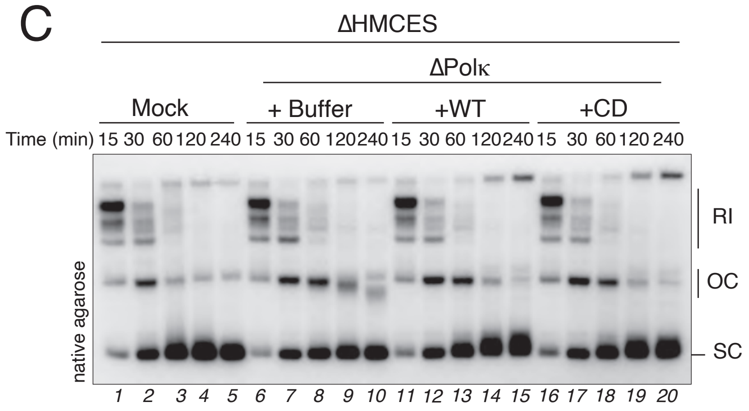

C
ΔHMCES
ΔPolκ
Mock
+ Buffer
+WT
+CD
Time (min) 15 30 60 120 240 15 30 60 120 240 15 30 60 120 240 15 30 60 120 240
native agarose
RI
OC
SC
1 2 3 4 5 6 7 8 9 10 11 12 13 14 15 16 17 18 19 20


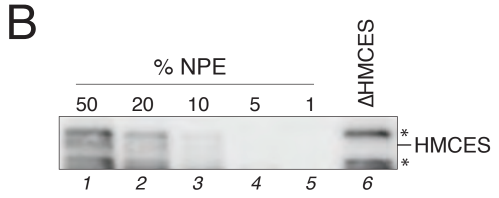

B
% NPE
50 20 10 5 1
ΔHMCES
*
HMCES
*
1 2 3 4 5 6

Figure S6

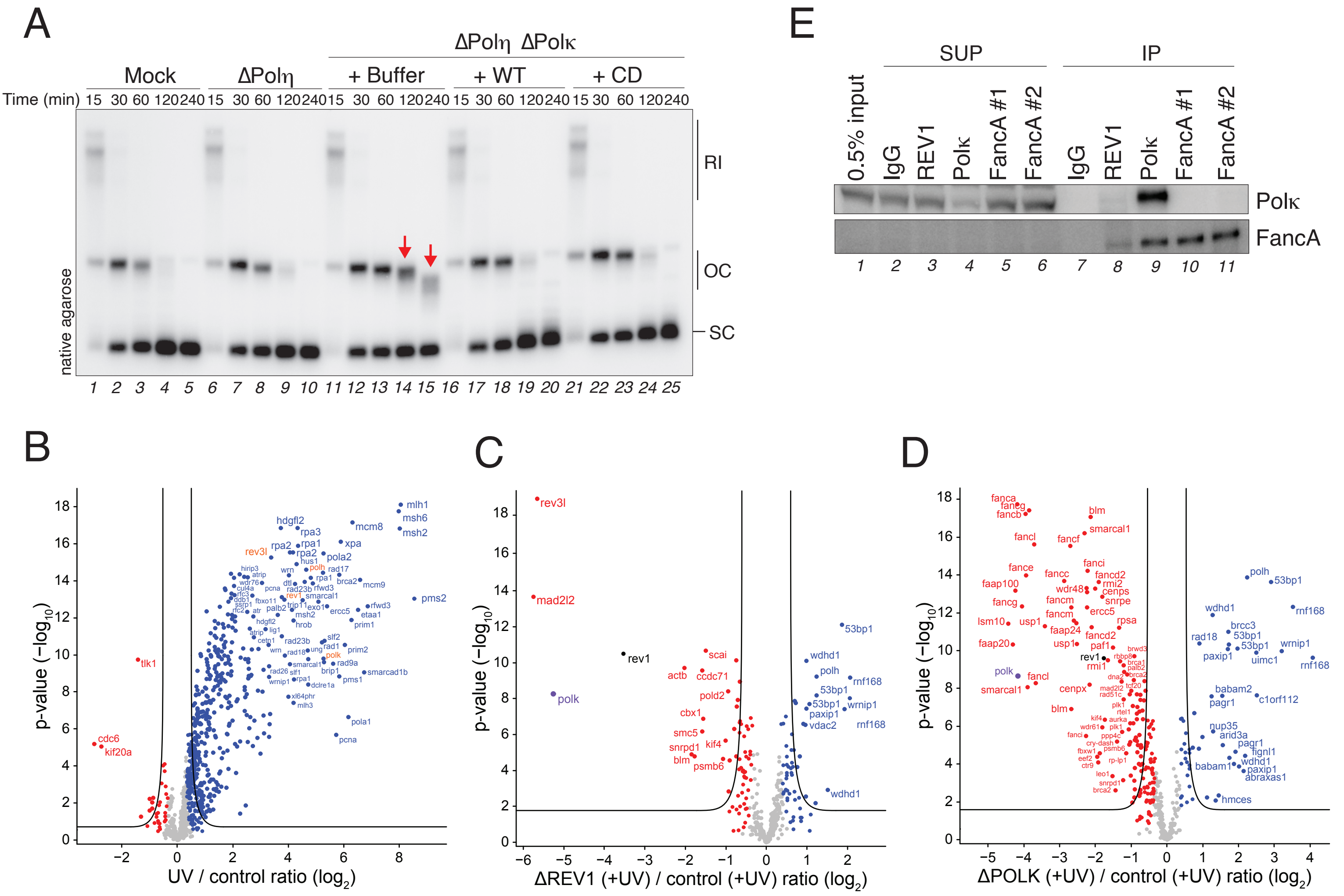
A
ΔPolη ΔPolκ
Mock
ΔPolη
+ Buffer
+ WT
+ CD
Time (min) 15 30 60 120 240 15 30 60 120 240 15 30 60 120 240 15 30 60 120 240 15 30 60 120 240
RI
OC
SC
native agarose
1 2 3 4 5 6 7 8 9 10 11 12 13 14 15 16 17 18 19 20 21 22 23 24 25
B
p-value (−log10)
UV / control ratio (log2)
mlh1
msh6
msh2
hdgfl2
rpa3
mcm8
rpa2
rpa1
xpa
rev3l
pola2
pms2
tlk1
cdc6
kif20a
pola1
pcna
C
p-value (−log10)
ΔREV1 (+UV) / control (+UV) ratio (log2)
rev3l
mad2l2
rev1
polk
scai
actb
ccdc71
pold2
cbx1
smc5
snrpd1
kif4
blm
psmb6
53bp1
wdhd1
polh
rnf168
wrnip1
paxip1
vdac2
D
p-value (−log10)
ΔPOLK (+UV) / control (+UV) ratio (log2)
fanca
fancg
fancb
blm
smarcal1
fancl
fancf
fanci
fance
fancc
fancd2
faap100
wdr48
rmi2
cenps
fancm
snrpe
lsm10
usp1
ercc5
faap20
faap24
rpsa
rev1
rmi1
polk
cenpx
polh
53bp1
rnf168
wdhd1
brcc3
rad18
paxip1
uimc1
wrnip1
babam2
pagr1
c1orf112
nup35
arid3a
fignl1
babam1
abraxas1
hmces
E
SUP
IP
0.5% input
IgG
REV1
Polκ
FancA #1
FancA #2
Polκ
FancA
1 2 3 4 5 6 7 8 9 10 11

A

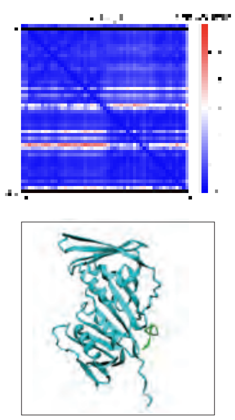

Q6DFE4_524-531;
P18248_R0
iptm: 0.915
iptm+ptm: 0.929

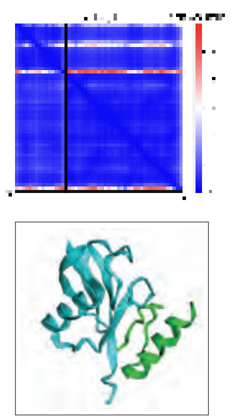

Q6DFE4_765-795;
P0CG47_1-76_R0
iptm: 0.791
iptm+ptm: 0.805

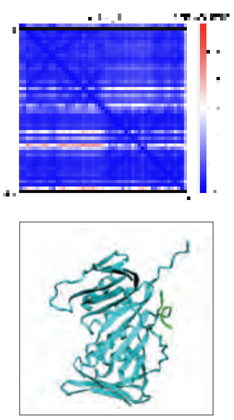

Q6DFE4_854-862;
P18248_R0
iptm: 0.915
iptm+ptm: 0.916

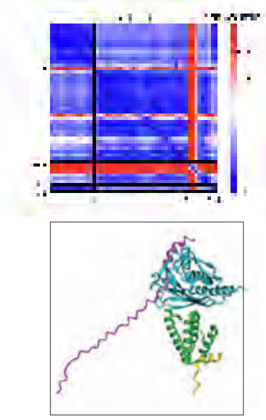

Q6NRK6_1136-1230;
Q8QFR4;
D0VEW8_1863-1913;
Q6DFE4_555-570_R0
iptm: 0.797
iptm+ptm: 0.796

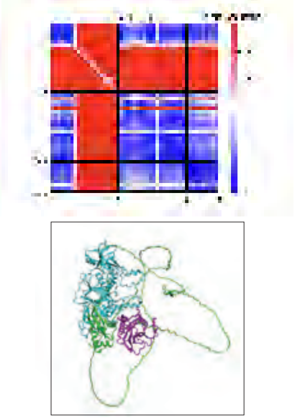

Q76LD3;
O93610;
Q8QFR4
iptm: 0.893
iptm+ptm: 0.856

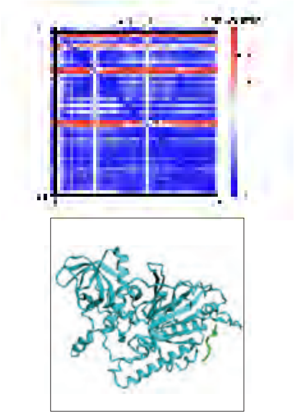

Q76LD3_132-142;
O93610_R0
iptm: 0.9213
iptm+ptm: 0.909

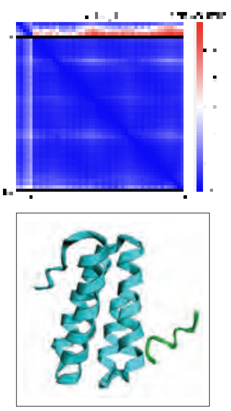

Q76LD3_232-239;
Q6NRK6_1146-1230_R0
iptm: 0.5447
iptm+ptm: 0.598

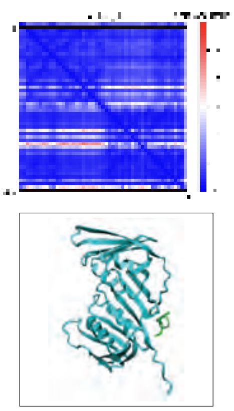

Q76LD3_444-452;
P18248_R0
iptm: 0.9175
iptm+ptm: 0.917

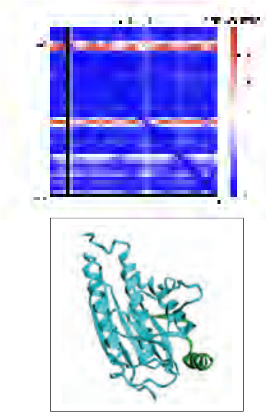

D0VEW8_2007-2028;
Q8QFR4_R0
iptm: 0.9479
iptm+ptm: 0.940

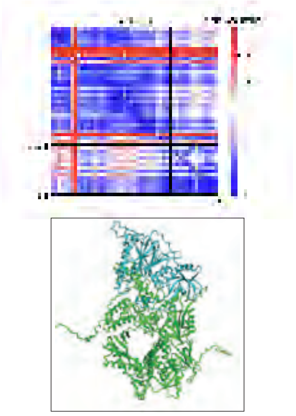

D0VEW8_1-200,
2232-3139;
O93610_R0
iptm: 0.9036
iptm+ptm: 0.895

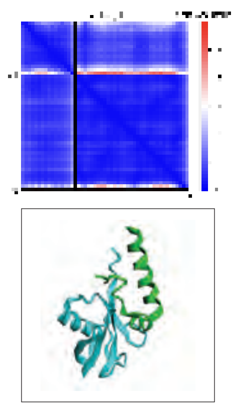

Q6NRK6_994-1027;
P0CG47_1-76_R0
iptm: 0.731
iptm+ptm: 0.749

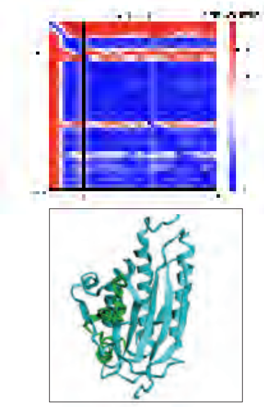

D0VEW8_1867-1917;
Q8QFR4_R0
iptm: 0.861
iptm+ptm: 0.857

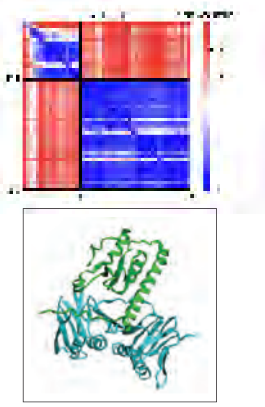

Q6NRK6_1-134;
P18248_R0
iptm: 0.188
iptm+ptm: 0.284

B

| PDB | Scaffold Structures |
|---|---|
| 7NV1 | *"Human Pol Kappa holoenzyme with Ub-PCNA"* |
| 3TBL | *"Structure of Mono-ubiquitinated PCNA"* |
| 6TNY | *"Processive human polymerase delta holoenzyme"* |
| 6V93 | *"Structure of DNA Polymerase Zeta/DNA/dNTP Ternary Complex"* |

| Single protein Prediction | plddt |
|---|---|
| Q6DFE4_1-520_R0 | 86.419 |
| Q6DFE4_1-531_R0 | 87.103 |
| Q8QFR4_R0 | 89.519 |
| Q6NRK6_R0 | 65.063 |
